# Chronic activation of a key exercise signal transducer, CaMKII, drives skeletal muscle aging and sarcopenia

**DOI:** 10.1101/2025.07.30.667744

**Authors:** Michael R. Bene, Tae Chung, William A. Fountain, Giovanni Rosales-Soto, Erick Hernández-Ochoa, Corina Antonescu, Liliana Florea, Seeun J. Jeong, Anne Le, Qian-Li Xue, Ahmet Hoke, Peter Abadir, Qinchuan Wang

**Affiliations:** Division of Geriatric Medicine and Gerontology, Johns Hopkins School of Medicine, Baltimore, Maryland, USA; Department of Physical Medicine and Rehabilitation, Johns Hopkins University School of Medicine, Baltimore, MD, USA; Department of Biochemistry and Molecular Biology, School of Medicine, University of Maryland, Baltimore, Maryland, USA; Genetic Resources Core Facility, Computational Bioanalysis Division, and Department of Genetic Medicine, Johns Hopkins School of Medicine, Baltimore, Maryland, USA; Gigantest, Inc., Baltimore, Maryland, USA; Departments of Neurology and Neuroscience, Johns Hopkins School of Medicine, Baltimore, MD, USA

## Abstract

Sarcopenia, the age-related loss of muscle strength and mass, contributes to adverse health outcomes in older adults. While exercise mitigates sarcopenia by transiently activating calcium (Ca^2+^)- and reactive oxygen species (ROS)-dependent signaling pathways that enhance muscle performance and adaptation, these same signals become chronically elevated in aged skeletal muscle and promote functional decline. Ca^2+^/calmodulin-dependent protein kinase II (CaMKII) is a key transducer of both Ca^2+^ and ROS signals during exercise. Here we show that CaMKII is chronically activated in aged muscles, promoting muscle dysfunction. Muscle-specific expression of a constitutively active CaMKII construct in young mice recapitulates features of aging muscles, including impaired contractility, progressive atrophy, mitochondrial disorganization, formation of tubular aggregates, and an older transcriptional profile characterized by the activation of inflammatory and stress response pathways. Mediation analysis identified altered heme metabolism as a potential mechanism of CaMKII-induced weakness, independent of muscle atrophy. Conversely, partial inhibition of CaMKII in aged muscle improved contractile function and shifted the transcriptome toward a more youthful state without inducing hypertrophy. These findings identify chronic CaMKII activation as a driver of functional and molecular muscle aging and support the concept that CaMKII exemplifies antagonistic pleiotropy, whereby its beneficial roles in promoting muscle performance and adaptation during youth may incur deleterious consequences in aging. We propose that persistent CaMKII activation in aged skeletal muscle reflects unresolved cellular stress and promotes maladaptive remodeling. Enhancing physiological reserve capacity through exercise, in combination with temporally targeted CaMKII inhibition, may help restore adaptive CaMKII signaling dynamics and preserve muscle function in aging.

## 1 INTRODUCTION

Skeletal muscle function peaks in young adulthood, followed by a plateau and then an accelerating decline in both muscle strength and mass from midlife onward (Dodds et al. 2014). In about 10% of community-dwelling older adults, this deterioration progresses to sarcopenia, a clinical disorder defined primarily by low muscle strength and secondarily by low muscle mass or quality (Cruz-Jentoft et al. 2019; Mayhew et al. 2019). Sarcopenia increases the risk of falls, disability, loss of independence, chronic diseases, and premature mortality, while placing a growing financial burden on healthcare systems through more frequent and costly hospitalizations (Cruz-Jentoft et al. 2019). These wide-ranging consequences highlight the need to delineate the molecular drivers of muscle aging and to identify tractable therapeutic targets.

Age-related decline in muscle function reflects the deterioration of mechanisms spanning multiple biological scales (Fielding et al. 2011; Tieland et al. 2018; Ferri et al. 2020). Given this complexity, exercise training remains the first-line intervention for both the prevention and treatment of sarcopenia, as it targets a broad array of biological processes implicated in muscle aging (Sayer & Cruz-Jentoft 2022). Exercise counteracts many features of muscle aging by activating an evolutionarily conserved adaptive program (Egan & Zierath 2013; Egan & Sharples 2023). Each bout of exercise disrupts skeletal muscle homeostasis (Winter & Fowler 2009), triggering signaling cascades that restore homeostasis and promote structural and functional remodeling. With repeated bouts, these adaptive responses enhance muscle performance and resilience.

Cytosolic calcium ions (Ca^2+^) not only control skeletal muscle force generation but also act as a signal for metabolic responses during muscle contraction and promote longer-term adaptations to exercise training (Berchtold et al. 2000). A key mediator of the Ca^2+^ signal is Ca^2+^/calmodulin-dependent protein kinase II (CaMKII) (Chin 2005; Bayer & Schulman 2019). CaMKII orchestrates diverse aspects of muscle physiology, regulating substrate uptake and utilization, Ca^2+^ cycling, and transcriptional programs involved in exercise adaptation (Flück et al. 2000; Liu et al. 2005; Raney & Turcotte 2008; Witczak et al. 2010; Ojuka et al. 2012; Ma et al. 2014; Cho et al. 2019; Wang et al. 2021; Flück et al. 2024).

Reactive oxygen species (ROS) constitute a second major signal in exercising muscles. While historically regarded as deleterious byproducts of metabolism, ROS generated during exercise are now recognized as essential signaling molecules that promote muscle adaptation (Powers et al. 2024). We have shown that CaMKII is activated by ROS in contracting skeletal muscle to augment both contractile performance and the transcriptional response to exercise (Wang et al. 2021).

Therefore, CaMKII mediates the physiological effects of Ca^2+^ and ROS in exercising skeletal muscle to promote performance and adaptation. In young, healthy muscle, the duration and degree of CaMKII activation are determined by the duration and intensity of contractile activity (Rose & Hargreaves 2003; Rose et al. 2006; Egan et al. 2010), as Ca^2+^ and ROS signals are regulated by muscle activity. However, aged skeletal muscle exhibits chronically elevated cytosolic Ca^2+^ and ROS at rest (Palomero et al. 2013; Mijares et al. 2021). This aberrant signaling environment likely drives persistent, exercise-independent activation of CaMKII. Given the broad impact of CaMKII on muscle physiology, we hypothesize that sustained, exercise-independent activation of CaMKII is detrimental, accelerating muscle aging and contributing to the pathogenesis of sarcopenia.

We first confirmed the upregulation and activation of CaMKII in the resting muscle of aged mice. We then used adeno-associated virus serotype 9 (AAV9) to deliver constitutively active CaMKIIγ to young muscle, modeling persistent activation while avoiding systemic confounders. We assessed its effects on muscle contractility, mass, histology, and transcriptomic profiles. Conversely, an AAV9-delivered CaMKII inhibitor was used to test the role of chronic CaMKII activation in muscle dysfunction in aged mice. Together, these experiments suggest chronic CaMKII activation as a driver of muscle aging and a potential target for intervention.

## 2 RESULTS

### CaMKII activity is elevated in skeletal muscle of older animals

To test the hypothesis that chronic activation of CaMKII contributes to muscle aging, we first evaluated CaMKII activity in skeletal muscle of sedentary older animals. We compared CaMKII autophosphorylation on threonine 287 (T287) (Brown & Bayer 2024), a marker for CaMKII activation, in tibialis anterior (TA) muscles of sedentary 33-month-old (old) and 3.7-month-old (young) mice using Western blot. The CaMKII isoforms, CaMKIIβ and CaMKIIδ/γ, exhibited significantly higher T287 phosphorylation in the old mice (Figure 1a and b), suggesting increased CaMKII activity in aged muscles without exercise.

**FIGURE 1.**
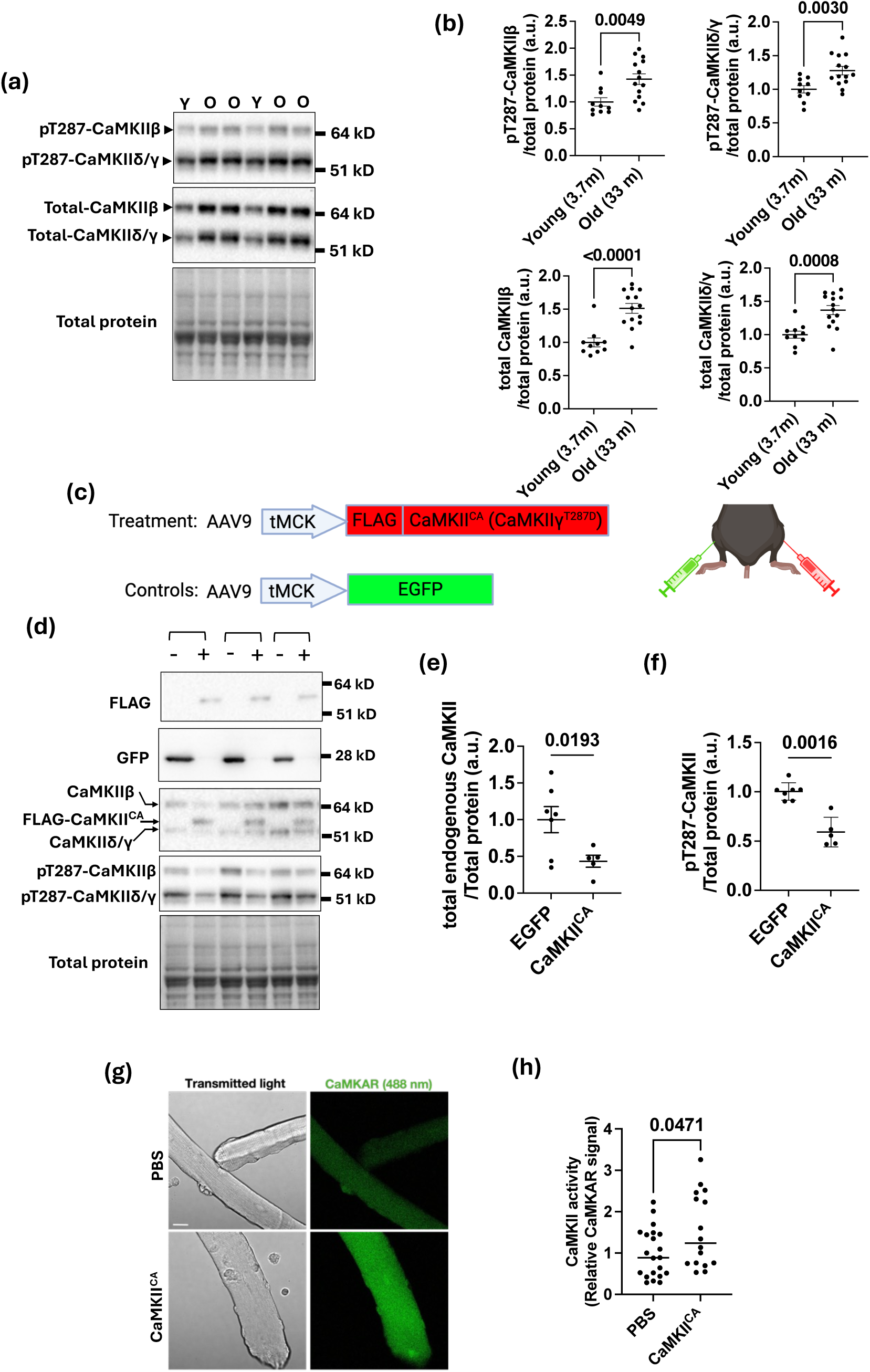
Age-associated elevation of CaMKII autophosphorylation and expression is recapitulated in young muscle by AAV9-mediated CaMKII^CA^ delivery. (a) Representative Western blots of CaMKIIβ, and CaMKIIδ/γ autophosphorylated on threonine 287 (pT287-CaMKII), total CaMKII, and total protein stain in the tibialis anterior (TA) muscles from young (Y, 3.7 months) and old (O, 33 months) mice. (b) Quantification of pT287-CaMKIIβ, pT287-CaMKIIδ/γ, total CaMKIIβ, and total CaMKIIδ/γ, normalized to total protein stain. n = 10 (young), and n = 14 (old); Student’s t-test, p-values in graphs. Data are mean ± SEM. (c) Schematic of AAV9 constructs for gene delivery. FLAG-tagged CaMKII^CA^ (CaMKIIγ-T287D) or Enhanced Green Fluorescent Protein (EGFP) were expressed under the muscle-specific tMCK promoter. AAV9 vectors were injected into TA muscles, with most mice receiving AAV9-tMCK-CaMKII^CA^ in one leg and AAV9-tMCK-EGFP in the contralateral leg for within-animal comparisons. (d) Western blot analysis of protein extracted from injected TA muscles. Each bracket indicates an individual mouse; (+) denotes AAV9-tMCK-CaMKII^CA^ injection, and (-) denotes AAV9-tMCK-EGFP injection. Blots were probed with anti-FLAG, anti-EGFP, anti-total-CaMKII, and anti-pT287-CaMKII antibodies. Total protein staining served as a loading control. (e, f) Quantification of endogenous total CaMKII and pT287-CaMKII levels, combining CaMKIIβ and CaMKIIδ/γ signals. n = 7 (EGFP), n = 5 (CaMKII^CA^); Student’s t-test, p-values shown in graphs. Data are mean ± SEM. (g) Confocal imaging of isolated single muscle fibers from flexor digitorum brevis (FDB) muscles expressing CaMKAR, a CaMKII activity reporter. Representative transmitted light and fluorescence (488 nm) images from PBS- and AAV9-tMCK-CaMKII^CA^-injected muscles are shown. (h) Quantification of CaMKII activity based on relative CaMKAR signal intensity in isolated fibers. n = 21 (PBS), n = 16 (CaMKII^CA^), with fibers isolated from three muscles per condition. Mann-Whitney test, p-value shown in the graph.

The total protein expression of these CaMKII isoforms was also significantly elevated in aged TA muscles, with the magnitude of upregulation comparable to that of T287 phosphorylation (Figure 1a, b). Similar increases of phosphorylated and total CaMKII were observed in the gastrocnemius muscles of 20-month-old mice compared to 3.7-month-old mice and in the quadriceps muscles of 30-month-old F344BN rats compared to 8-month-old rats (data not shown), consistent with previous reports (Chin 2004; Ljubicic & Hood 2009; Kramerova et al. 2012). Together, these findings suggest that CaMKII protein expression is upregulated in aged muscles. Furthermore, the intracellular milieu of aged muscles sustains CaMKII autophosphorylation even in the absence of exercise.

### Adeno-associated virus serotype 9-mediated expression of constitutively active CaMKII increases CaMKII activity in skeletal muscle

To model the chronically elevated expression and autophosphorylation of CaMKII in aged muscles, we used AAV9 to express FLAG-tagged CaMKIIγ carrying a phosphomimetic T287D mutation that renders it constitutively active (CaMKII^CA^) (Figure 1c). The CaMKII^CA^ was controlled by the muscle-specific tMCK promoter, confining its expression to muscle fibers (Wang et al. 2008). AAV9 was selected for its low immunogenicity and efficient transgene delivery in mouse skeletal muscle (Muraine et al. 2020). The CaMKII^CA^ construct originated from CaMKIIγ transcript variant 3 (NM_001039139), the most abundant CaMKII isoform in skeletal muscle, as determined by our previously published transcriptomic data (Wang et al. 2021). The AAV9-tMCK-CaMKII^CA^ was delivered via intramuscular injection into one TA muscle per mouse. Concurrently, the contralateral TA was injected with an equal viral load of AAV9-tMCK-Enhanced Green Fluorescent Protein (EGFP) as a control. This contralateral design enables paired comparisons of the TA muscles within individual animals, thereby minimizing influence by inter-individual variabilities.

Western blotting confirmed the expression of the FLAG-tagged CaMKII^CA^ and EGFP within the TAs, with very little ectopic expression detected in respective contralateral muscles, indicating that transgene expression remained localized to the injected muscles (Figure 1d). As anticipated, CaMKII^CA^ was detected using the anti-total CaMKII antibody, and its expression level was comparable to endogenous CaMKII (Figure 1d). The pT287-CaMKII antibody did not detect CaMKII^CA^, confirming its specificity, as the T287D mutation eliminates T287 phosphorylation. Notably, the expression and T287 phosphorylation of endogenous CaMKII were significantly reduced in CaMKII^CA^-expressing muscles (Figure 1d, e, and f), suggesting potential negative feedback mechanisms that limit CaMKII protein levels and activity. This phenomenon raised the question whether net CaMKII activity was indeed elevated in CaMKII^CA^-expressing muscles.

We therefore directly measured CaMKII activity in single muscle fibers isolated from the Digitorum Brevis (FDB) muscles. For these measurements, in each mouse, one FDB muscle was injected with AAV9-tMCK-CaMKII^CA^, while the contralateral FDB, serving as the control, was injected with an equal volume of phosphate-buffered saline (PBS). All muscles were subsequently electroporated with a plasmid encoding the CaMKII activity reporter, CaMKAR (Reyes Gaido et al. 2023; Severino et al. 2025), allowing live-cell CaMKII activity measurement via confocal imaging (Figure 1g). PBS instead of AAV9-tMCK-EGFP was used as the control specifically for this fluorescence-based assay to avoid spectral overlap from EGFP, as CaMKAR is a GFP-derived ratiometric reporter. The CaMKAR fluorescent signal was significantly higher in the CaMKII^CA^-expressing fibers than in control fibers (Figure 1h). These findings demonstrate that despite reducing total and T287 phosphorylated endogenous CaMKII, CaMKII^CA^ expression effectively raises net CaMKII activity in muscle fibers. This validates AAV9-tMCK-CaMKII^CA^ as a suitable tool for investigating the consequences of chronically elevated CaMKII activity in skeletal muscles.

### Sustained CaMKII activation progressively weakens tibialis anterior muscles and induces histological remodeling

Having confirmed elevated CaMKII activity in aged skeletal muscle and validated that AAV9-tMCK-CaMKII^CA^ can recapitulate this activation in young muscle, we next assessed the functional impact of sustained CaMKII activation in vivo. To isolate these effects from age-related confounders, we injected 3.5-month-old mice with AAV9-tMCK-CaMKII^CA^ in one TA and the AAV9-tMCK-EGFP in the contralateral TA. Six weeks later, in vivo contractility was assessed by stimulating the peroneal nerve across a frequency range of 1–150 Hz in anesthetized mice. CaMKII^CA^-expressing muscles produced significantly less force than their EGFP-expressing counterparts across all frequencies (estimate = -1.13 [-1.44, -0.82], p = 1.44 × 10^-11^, Figure 2a). At the seven-week endpoint, CaMKII^CA^-expressing TAs weighed about 7.0% less than paired controls (p < 0.0001, Figure 2b), suggesting muscle atrophy. When the measured forces were normalized against the muscle mass, the normalized force remained significantly smaller in CaMKII^CA^-expressing muscles (estimate = -1.87 [-2.47, -1.28], p = 1.07 × 10^-4^, Figure 2c). Mediation analysis showed a significant direct effect of CaMKII^CA^ on contractility (Average Direct Effect [ADE]: -1.10, p ≤ 2 × 10^-16^, Figure 2d), while muscle mass did not significantly mediate this effect (Average Causal Mediation Effect [ACME]: -0.03, p = 0.82). Consequently, the Total Effect of CaMKII^CA^ on force (-1.13, p ≤ 2 × 10^-16^) was almost entirely attributed to its direct negative influence. Thus, sustained CaMKII activation for 6–7 weeks compromises muscle contractile performance by reducing the intrinsic force-generating capacity while also inducing modest atrophy.

**FIGURE 2.**
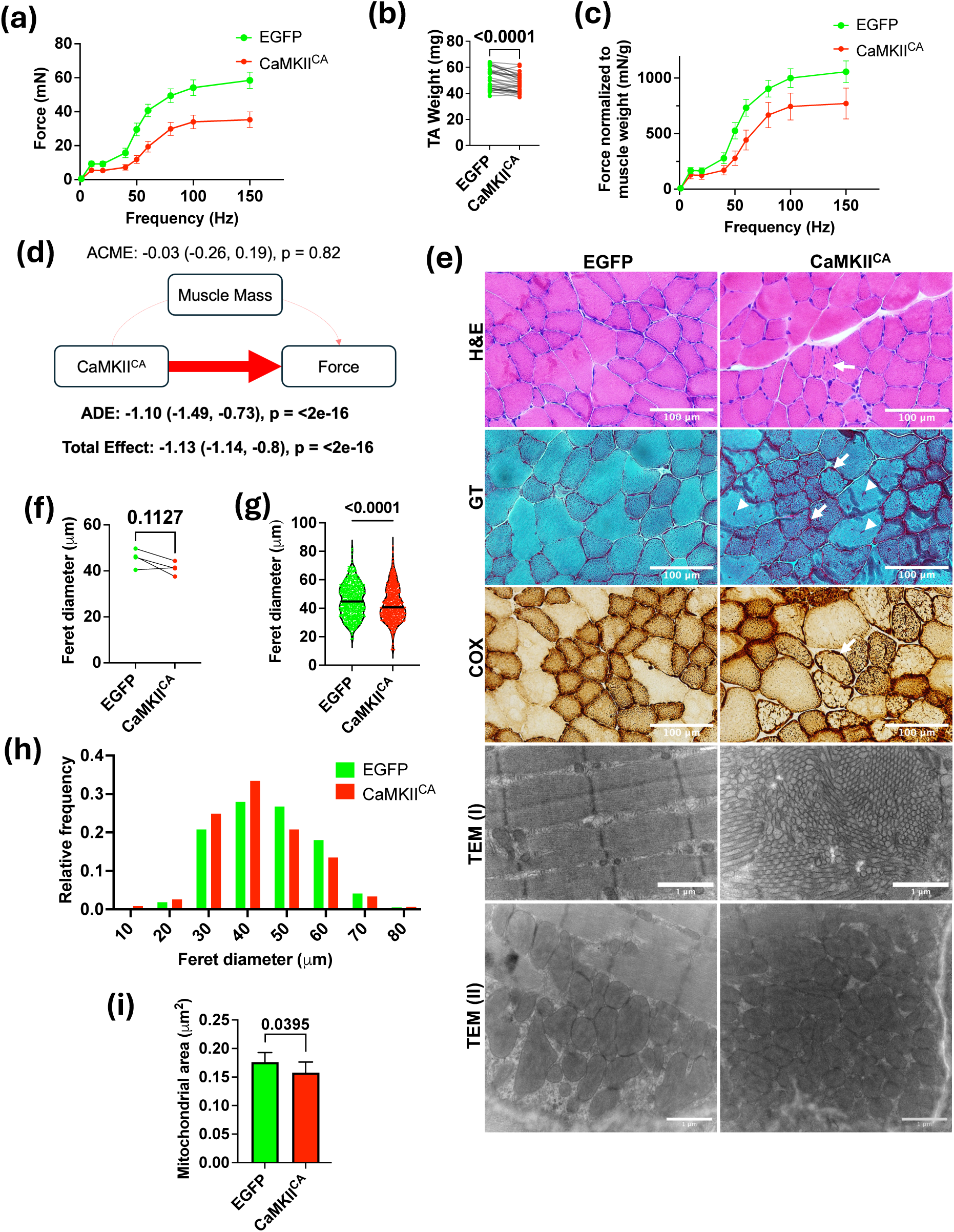
CaMKII activation for 6 weeks impairs muscle contractility and induces atrophy and histological changes. (a) In vivo isometric force-frequency curves of TA muscles 6 weeks post-injection of AAV9-tMCK-EGFP (one TA) or AAV9-tMCK-CaMKII^CA^ (contralateral TA). n = 9 per condition; linear mixed-effects model, estimate = -1.13 (-1.44,-0.82), p = 1.44 × 10^-11^, p-value in graph. (b) TA muscle mass 7 weeks post-injection. n = 30 pairs; paired Student’s t-test, p-value in graph. (c) Isometric force normalized to TA mass. n = 10 per condition; linear mixed-effects model, estimate = -1.87 [-2.47, -1.28], p = 1.07 × 10^-4^. (d) Mediation analysis (“mediation” R package). (e) Representative H&E, modified Gomori’s Trichrome (GT) staining, and COX staining of TA cross-sections 7 weeks post-injection, as well as two sets of transmission electron microscopy (TEM) images from comparable regions of the muscles. (f) Muscle fiber minimum Feret diameters. Each point represents the median of ∼200 fibers per muscle. n = 4 muscles per treatment; paired Student’s t-test, p-value in graph. (g) Muscle fiber minimum Feret diameters, pooled by treatment. n = 755 (EGFP), n = 808 (CaMKII^CA^). (h) Histogram of relative frequency distributions of muscle fiber minimum Feret diameters. n = 755 (EGFP), n = 808 (CaMKII^CA^). (i) Quantification of mitochondrial areas. n = 720 (EGFP), n = 582 (CaMKII^CA^), TEM images were acquired from 4 muscles for each treatment. The bar graph shows the median and 95% confidence interval; Mann-Whitney test, p-value in the graph.

Given that by mediation analysis, reduced muscle mass did not account for the CaMKII^CA^-induced reduction in contractile force, we conducted histological analyses 7 weeks post-injection to examine additional structural changes associated with impaired function (Figure 2e – i). Routine H&E staining revealed an absence of endomysial inflammation or necrotic fibers in CaMKII^CA^-expressing TA muscles. Morphometric analysis of H&E-stained muscle sections from four mice, each expressing CaMKII^CA^ in one TA and EGFP in the contralateral TA, revealed a non-significant trend toward smaller minimum Feret diameters of muscle fibers in CaMKII^CA^-expressing muscles (∼200 fibers per muscle from the superficial and deep regions per section, paired t-test, p = 0.1127; Figure 2f). However, pooling all measurements by treatments showed that CaMKII^CA^-expressing muscles had significantly smaller minimum Feret diameters (p < 0.0001, Figure 2g), and a leftward shift in the diameter distribution (Figure 2h), confirming the presence of muscle fiber atrophy.

Some CaMKII^CA^-expressing muscle fibers exhibited an altered basophilic staining pattern in H&E-stained sections (Figure 2e, H&E panel, arrow). Modified Gomori’s trichrome (GT) staining revealed features of tubular aggregates (Figure 2e, GT panel, arrowheads), which are structures formed by sarcoplasmic reticulum (SR) aggregation and are frequently found in aged skeletal muscle and muscles with defective Ca^2+^ regulation (Boncompagni et al. 2021). The presence of tubular aggregates in CaMKII^CA^-expressing muscle was confirmed by transmission electron microscopy (Figure 2e, TEM panel I).

GT staining revealed some muscle fibers containing dense subsarcolemmal red deposits (Figure 2e, GT panel, arrow), reminiscent of features of ragged-red fibers caused by mitochondrial dysfunction (Rifai et al. 1995; Dubowitz et al. 2020). However, histochemical staining for cytochrome c oxidase (COX) and succinate dehydrogenase (SDH) revealed a redistribution of these mitochondrial enzymatic activities toward subsarcolemmal regions in affected fibers, with preservation of both COX and SDH activities (Figure 2e, COX panel, arrow; SDH staining not shown). The preservation of COX activity distinguishes these fibers from ragged-red fibers resulting from mitochondrial DNA mutations, where COX activity is impaired. TEM images showed that CaMKII^CA^-expressing muscles had prominent clumps of mitochondria in the subsarcolemmal region (Figure 2e TEM panel II), which is consistent with findings from COX staining. Furthermore, CaMKII^CA^-expressing muscles had smaller mitochondria (Figure 2i).

To assess the long-term consequences of sustained CaMKII activation, a separate cohort of mice injected with AAV9-tMCK-CaMKII^CA^ in one TA and PBS in the contralateral TA at 3.5 months of age was evaluated 9 months post-injection. This cohort was established before the availability of the EGFP control virus. As prior work has shown that AAV9-EGFP exerts minimal effects on skeletal muscle when compared with PBS injections (Riaz et al. 2015), PBS serves as a valid control for evaluating CaMKII^CA^-specific effects. Nine months after injection, CaMKII^CA^-expressing muscles exhibited impaired contractile force compared to the PBS-injected control (estimate = -0.99 [-1.39,-0.59], p = 4.21 × 10^-6^, Supplementary Figure 1a). Reduction in muscle mass progressed from 7.0% at 7 weeks after CaMKII^CA^ expression to 24.2% after 9 months (Supplementary Figure 1b). Contractile force of CaMKII^CA^-expressing muscles remained significantly smaller after normalization against muscle mass (estimate = -1.25 [-1.87,-0.62], p = 1.48 × 10^-4^, Supplementary Figure 1c). However, in contrast to the 6-week timepoint, when force loss was not attributed to atrophy, mediation analysis at 9 months showed that the decline in contractile force was explained by muscle mass loss (ACME: -1.16, p ≤ 2 × 10^-16^, Supplementary Figure 1d).

Building on the structural alterations observed at 7 weeks and considering the more pronounced functional decline and atrophy at 9 months, we examined muscle histology to evaluate potential progression of tissue remodeling (Supplementary Figure 1e–h). H&E staining revealed significant numbers of atrophic fibers in CaMKII^CA^-expressing muscles, and the absence of endomysial inflammation or necrotic fibers (Supplementary Figure 1e, H&E panel). Morphometric analysis revealed a significant reduction in median minimum Feret diameters in CaMKII^CA^ muscles compared to contralateral controls (p = 0.0143; Supplementary Figure 1f). Pooling all measurements by treatments also showed that CaMKII^CA^-expressing muscles had significantly smaller median fiber diameters (p<0.0001, Supplementary Figure 1g), and a leftward shift in the diameter distribution (Figure 2h)

GT staining showed further changes in staining patterns compared to control sections (Supplementary Figure 1e, GT panel), suggesting further remodeling of the sarcoplasmic reticulum and mitochondria. COX staining showed an extensively disorganized mitochondrial network with preserved COX activity (Supplementary Figure 1e, COX panel). Despite the extensive remodeling of the mitochondrial network, CaMKII^CA^ did not measurably alter mitochondrial protein content relative to total muscle protein. Immunoblotting for representative subunits of respiratory-chain complexes I–V revealed no significant difference in their steady-state abundance at either 7 weeks (Supplementary Figure 2a and b) or 9 months (Supplementary Figure 3a and b) after viral injection. This included nuclear-encoded subunits (NDUFB8, SDHB, UQCRC2, ATP5A) and the mitochondrial-encoded subunit MTCO1, indicating preserved stoichiometric balance between nuclear and mitochondrial genome-encoded components of the respiratory chain complexes. Consistently, quantitative PCR showed an unchanged mitochondrial-to-nuclear DNA copy number ratio at 7 weeks (Supplementary Figure 2c). These findings suggest that CaMKII^CA^ alters mitochondrial spatial organization and fission/fusion dynamics without affecting the abundance of mitochondrial constituents in skeletal muscle.

### Chronic CaMKII activation shifts the transcriptome of young muscle toward an aged profile, whereas partial CaMKII inhibition modestly rejuvenates that of aged muscle

Sustained CaMKII activation impairs muscle function, alters fiber morphology, and disrupts mitochondrial and SR organization, suggesting that chronic CaMKII signaling may also drive transcriptional remodeling. To test this, we performed poly(A) RNA sequencing (RNA-seq) on TA muscles seven weeks after in vivo modulation of CaMKII activity (Figure 3a). In young adult mice (3.5 months; 4 males, 5 females), one TA was injected with AAV9-tMCK-CaMKII^CA^ and the contralateral TA with AAV9-tMCK-EGFP, enabling within-subject comparisons between CaMKII^CA^-expressing and control muscles, independent of aging-related confounders. The seven-week time point was chosen to capture transcriptomic changes associated with early functional and histological alterations, before the extensive structural degeneration observed at later stages. In parallel, aged mice (21.5 months; 4 males, 4 females) were injected with AAV9-tMCK-CN19o in one TA and AAV9-tMCK-EGFP in the contralateral TA to assess the effects of CaMKII inhibition in an aged context by CN19o, a well-characterized peptide inhibitor of CaMKII (Coultrap & Bayer 2011).

**FIGURE 3.**
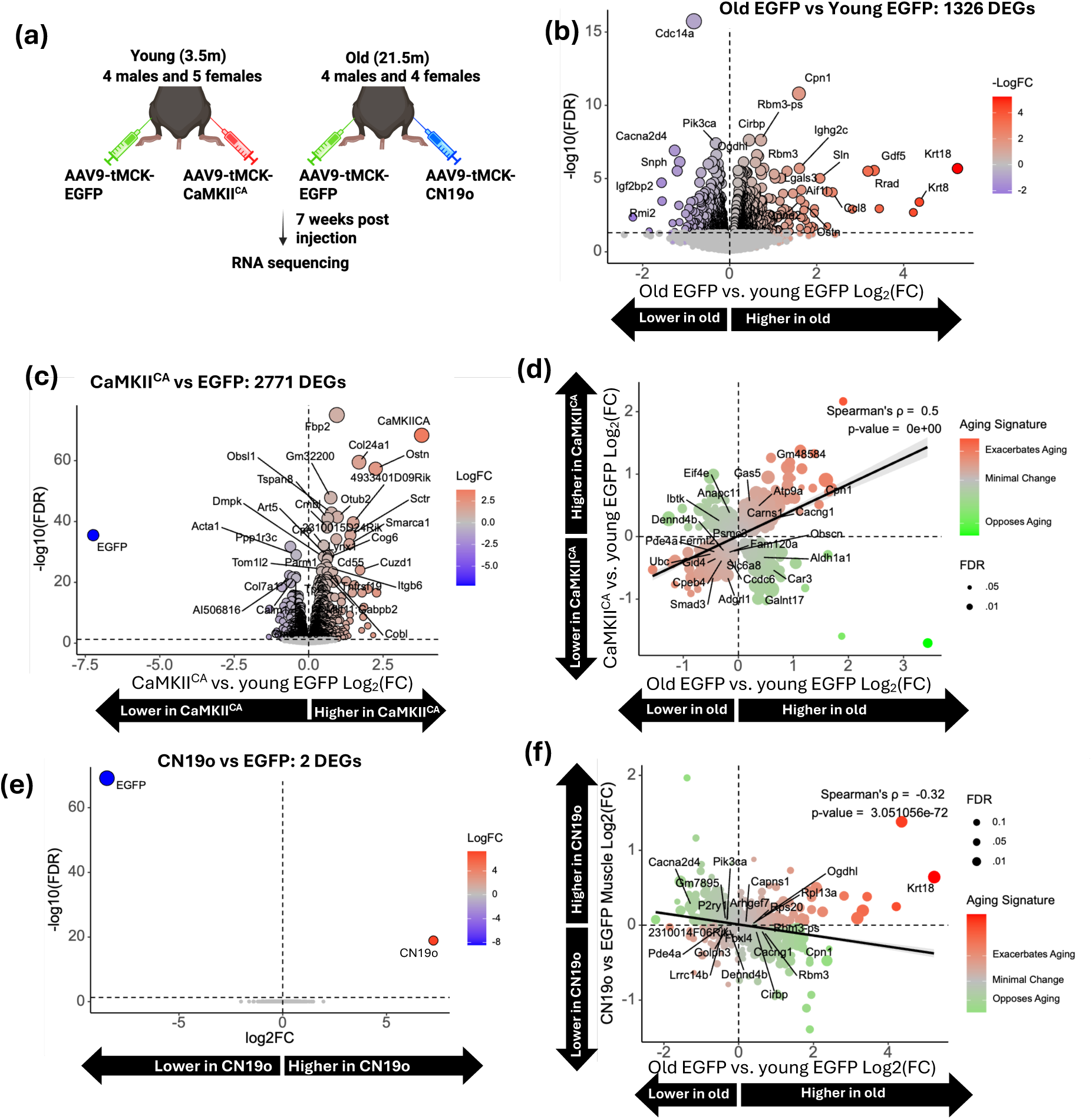
Constitutively active CaMKII (CaMKII^CA^) shifts the young muscle transcriptome toward an older profile, while CaMKII inhibition by CN19o partially rejuvenates the transcriptome of aged muscles. (a) Experimental design. Young mice (3.5m, n = 9, 4 males and 5 females) were injected with AAV9-tMCK-EGFP in one TA and AAV9-tMCK-CaMKII^CA^ in the contralateral TA. Old mice (21.5 months, n = 8, 4 males and 4 females) were injected with AAV9-tMCK-EGFP in one TA and AAV9-tMCK-CN19o (CaMKII inhibitory peptide) in the contralateral TA. TA muscles were collected 7 weeks post-injection and subjected to RNA sequencing. (b) Volcano plot comparing gene expression between old and young EGFP-expressing TA muscles; differentially expressed genes (DEGs). (c) Volcano plot comparing gene expression between young CaMKII^CA^- and young EGFP-expressing TA muscles. (d) Correlation analysis between genes regulated by aging (old vs. young EGFP) and by CaMKII^CA^ expression (young CaMKII^CA^ vs. young EGFP). Color indicates whether CaMKII^CA^ promoted, opposed, or minimally influenced aging-like changes of gene expression. Spearman’s ρ and p-value are shown. A linear trendline was included to indicate the overall correlation. (e) Volcano plot comparing gene expression between old CN19o-and old EGFP-expressing TA muscles. (f) Correlation analysis between genes regulated by aging (old vs. young EGFP) and by CN19o expression (old CN19o vs. old EGFP). Color indicates whether CN19o promoted, opposed, or influenced aging-like changes of gene expression. Spearman’s ρ and p-value are shown. A linear trendline was included to indicate the overall correlation.

Differential expression analysis identified 1,326 differentially expressed genes (DEGs) between old and young EGFP-expressing TAs (Figure 3b), consistent with the extensive transcriptional remodeling previously reported in aging rodent skeletal muscles (Graber et al. 2023). Seven weeks of CaMKII^CA^ expression in young muscle elicited an even broader response, with 2,771 genes differentially expressed relative to contralateral EGFP controls (Figure 3c). To determine how closely CaMKII^CA^ expression recapitulates the aging transcriptional program, we correlated the Log_2_(fold changes) for all genes in the two comparisons (young CaMKII^CA^ vs. young EGFP and old EGFP vs. young EGFP). The correlation showed a moderate but highly significant concordance (Spearman’s ρ = 0.50, p ≈ 0; Figure 3d). Of the 1,326 genes significantly altered with age, 73.5% were regulated in the same direction by CaMKII^CA^, while 26.5% showed discordant regulation. Conversely, 69.9% of the 2,771 CaMKII^CA^-regulated genes were shifted in the same direction by aging, while 31.1% were discordant. These data indicate that chronic CaMKII activation recapitulates a substantial portion of the aging-associated transcriptional changes while also driving a distinct set of gene changes, suggesting that additional pathways cooperate with CaMKII or act independently to shape the full breadth of age-related transcriptomic remodeling.

In aged TA muscles, AAV-driven expression of the CaMKII inhibitor CN19o resulted in only a minimal number of statistically significant transcriptional changes: after false-discovery correction (FDR q < 0.05), only the two transgene mRNAs, CN19o and EGFP, met the differential expression threshold (Figure 3e). This muted response is likely attributable to the low transcript abundance of CN19o relative to endogenous CaMKII isoforms (mean transcript length-normalized counts ± SEM: CN19o 0.56 ± 0.15; Camk2b 1.40 ± 0.13; Camk2d 0.22 ± 0.03; Camk2g 0.89 ± 0.05), suggesting that CN19o expression may have achieved only partial inhibition of CaMKII enzymatic activity, thereby limiting its transcriptional impact.

Although few individual genes crossed the FDR threshold, CN19o nonetheless resulted in a global expression pattern that opposed the transcriptional changes observed with aging. Spearman’s correlation of Log_2_(fold changes) for all genes revealed a significant inverse relationship between CN19o- and aging-associated expression changes (ρ = -0.32, p-value = 3.05×10^-72^; Figure 3f). This broad but subtle opposing effect suggests that even partial inhibition of CaMKII activity can counteract age-associated transcriptomic drift through coordinated, low-magnitude modulation of many genes.

### Sustained CaMKII activation recapitulates aging-associated pathway signatures

To understand the transcriptomic shifts, we performed gene set enrichment analysis (GSEA) using the Hallmark collection, which comprises 50 curated, non-redundant gene sets representing major biological processes (Liberzon et al. 2015; Subramanian et al. 2005). The analysis was performed on three comparisons: old EGFP vs young EGFP, young CaMKII^CA^ vs young EGFP, and old CN19o vs old EGFP. The resulting normalized enrichment scores (NES) were compared side-by-side (Figure 4a).

**FIGURE 4.**
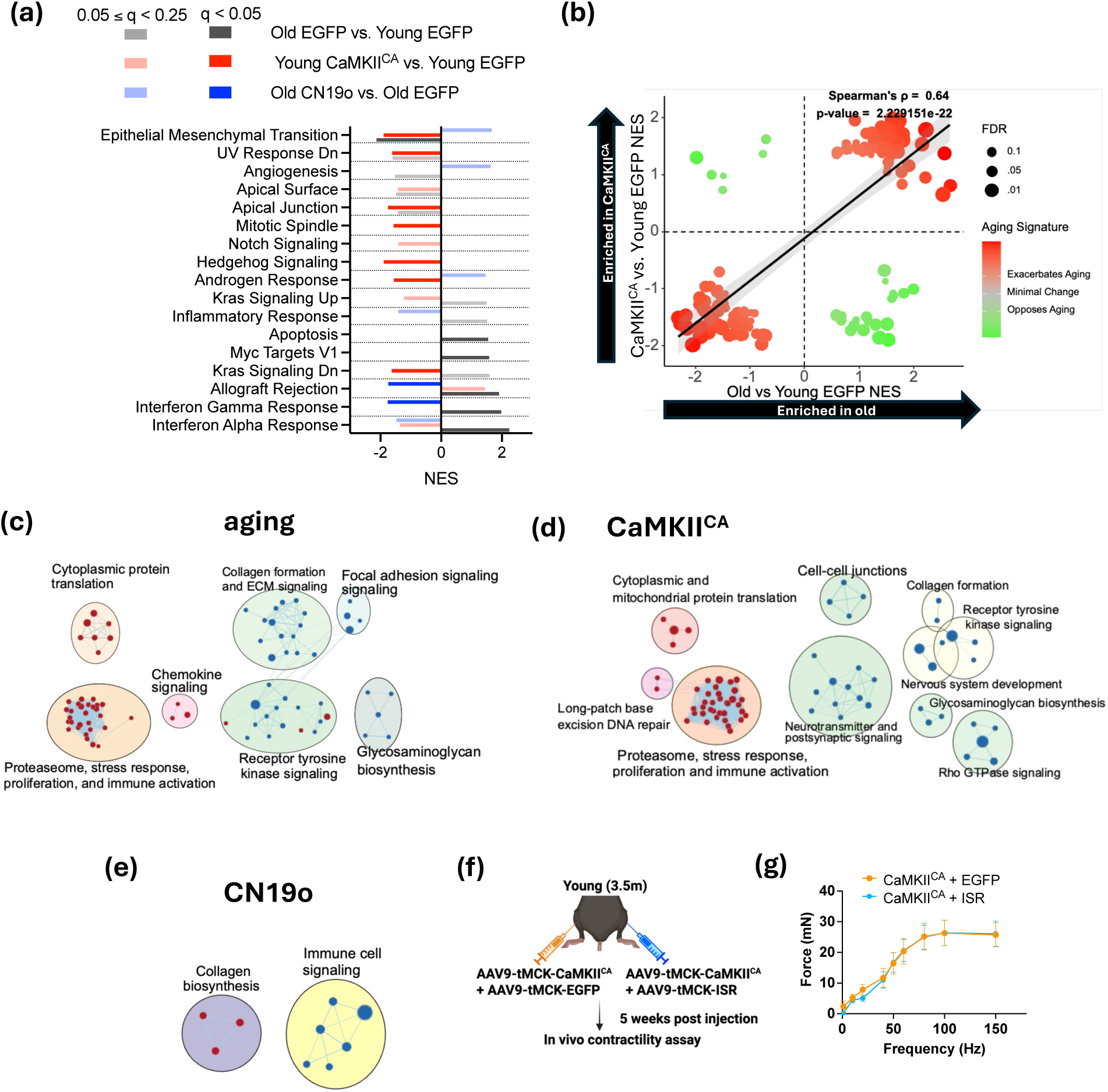
CaMKII activation drives aging-related pathway changes; its inhibition reverses inflammatory signatures in aged muscle, but NF-κB blockade fails to rescue CaMKII^CA^-induced contractile dysfunction. (a) Normalized Enrichment Scores (NES) from Gene Set Enrichment Analysis (GSEA) on indicated ranked gene lists. The FDR q-value ranges of the enriched pathways are indicated by color. (b) Correlation analysis between the NES of Reactome gene sets regulated by aging (old EGFP vs. young EGFP) or by CaMKII^CA^ (young CaMKII^CA^ vs. young EGFP). Color indicates whether CaMKII^CA^ exacerbated, opposed, or minimally influenced aging-related changes. Spearman’s ρ and p-value are shown. A linear trendline was included to indicate the overall correlation. (c and d) EnrichmentMap clustering of Reactome gene sets regulated by aging or by CaMKII^CA^; red and blue circles indicate positively and negatively enriched pathways, respectively; edges indicate overlap in underlying genes between connected pathways. (e) EnrichmentMap clustering of Reactome gene sets regulated by CN19o in old muscles compared to old EGFP expressing muscles. (f) Schematics of experimental design testing the effect of IκBα Super Repressor (ISR) on CaMKII^CA^-induced muscle weakness. (g) In vivo isometric force-frequency curves of TA muscles 5 weeks post-injection of AAV9-tMCK-CaMKII^CA^ with co-injection of either AAV9-tMCK-EGFP or AAV9-tMCK-ISR. n = 9 per condition; p-value in graph (p = 0.1).

GSEA revealed that sustained CaMKII activation modulates many of the same pathways altered in aging, while CaMKII inhibition in aged muscle reverses several of these changes. Among the 17 Hallmark gene sets meeting a relaxed significance threshold (FDR < 0.25 in any comparison), several showed similar enrichment patterns in both CaMKII^CA^-expressing young muscle and aged muscle (old EGFP). Gene sets negatively enriched in both CaMKII^CA^-expressing and aged muscle included Epithelial-Mesenchymal Transition, UV-Response Dn, Apical Surface, Apical Junction, and Interferon-α Response. Conversely, Allograft Rejection was positively enriched in both conditions. CN19o expression in aged muscle generally reversed these patterns. For example, Epithelial-Mesenchymal Transition, Angiogenesis, and Androgen Response shifted from negative to positive enrichment, while inflammatory signatures, including Inflammatory Response, Allograft Rejection, and Interferon-γ and -α Responses, shifted from positive to negative enrichment.

Some pathways exhibited discordant regulation between aging and CaMKII activation. For example, K-RAS Signaling and Interferon-α Response were enriched in opposite directions in the two conditions. Additionally, several pathways were selectively altered by either aging or CaMKII^CA^ expression but not both. These findings indicate that while sustained CaMKII activation recapitulates a substantial portion of the aging-associated pathway profile, other transcriptional programs in aging are likely driven by CaMKII-independent mechanisms.

To gain more detailed insight into pathway-level changes, we performed GSEA using gene sets from the Reactome Canonical Pathways, which offer finer granularity than the Hallmark gene sets that represent broad biological processes. Of 1,289 Reactome gene sets tested, 184 met the FDR < 0.25 threshold in either young CaMKII^CA^ vs. young EGFP or old EGFP vs. young EGFP comparisons (Supplementary Table 1). Normalized enrichment scores for young CaMKII^CA^ and οld EGFP conditions were significantly correlated (Spearman ρ = 0.64, p = 2.2 × 10^-22^; Figure 4b), indicating that chronic CaMKII activation recapitulates many aging-associated pathway changes. EnrichmentMap clustering (Reimand et al. 2019) revealed shared activation of proteasomal activity, cellular stress responses, immune signaling, and cytoplasmic protein translation (Figure 4c and d), consistent with accelerated protein turnover and a pro-inflammatory transcriptional environment. Both conditions also downregulated pathways related to extracellular matrix organization, including collagen synthesis, glycosaminoglycan biosynthesis, and receptor tyrosine kinase signaling, suggesting a shared suppression of anabolic and structural remodeling programs.

Despite the broad overlap with aging-associated pathways, CaMKII^CA^ induced distinct transcriptomic changes not observed in aged muscle. Notably, mitochondrial protein translation and long-patch base-excision DNA repair pathways were uniquely enriched, suggesting a heightened mitochondrial stress response and increased oxidative DNA damage. CaMKII^CA^ additionally down-regulated clusters linked to neurotransmitter and synaptic signaling, and Rho GTPase-mediated signaling, alterations that may underlie the early decline in contractile performance observed after seven weeks of expression.

Consistent with its limited effect at the single-gene level, CaMKII inhibition by CN19o in aged muscle produced few pathway-level changes. Only two Reactome pathway clusters, immune-cell signaling and collagen biosynthesis, met the FDR threshold, and both were regulated in the opposite direction to their enrichment in aging and CaMKII^CA^-expressing muscle (Figure 4e). These findings suggest that even partial CaMKII inhibition can suppress age-associated inflammatory and extracellular matrix remodeling programs via modest, coordinated transcriptional shifts.

Together, GSEA across Hallmark and Reactome gene sets demonstrates that aging and chronic CaMKII activation elicit broadly similar transcriptional changes, including upregulation of immune and inflammatory pathways. In contrast, CaMKII inhibition by CN19o selectively suppresses these immune-related signatures with limited effects on other pathways. These results support the concept that sustained CaMKII activity promotes a pro-inflammatory transcriptional state in skeletal muscle, consistent with our previous observation that CaMKII activation contributes to the transient induction of inflammatory genes following acute exercise (Wang et al. 2021).

Given that NF-κB is a central driver of chronic inflammation in aging tissues (Adler et al. 2007), is activated by CaMKII in various cell types (Singh et al. 2009; Martin et al. 2018; Yao et al. 2022), and promotes muscle wasting when hyperactivated (Cai et al., 2004), we tested whether canonical NF-κB signaling mediates the functional decline induced by CaMKII^CA^. We injected AAV9-tMCK-CaMKII^CA^ into both TAs of 3.5-month-old mice, co-injected AAV9-tMCK-IκBα-super-repressor (ISR) into one TA, and AAV9-tMCK-EGFP into the contralateral TA as control (Figure 4f). ISR has been shown to inhibit NF-κB signaling and attenuate muscle wasting in models of denervation and cancer cachexia (Cai et al. 2004). However, in vivo contractility assays performed five weeks post-injection revealed no improvement in force generation in ISR-treated muscles (Figure 4g). These results suggest that the canonical NF-κB pathway is not required for CaMKII^CA^-induced muscle weakness, indicating that other CaMKII-dependent mechanisms underlie the observed functional decline.

### The heme metabolism gene set enrichment score is a principal transcriptomic correlate of CaMKII^CA^-induced muscle weakness

To identify transcriptional mechanisms underlying CaMKII^CA^-driven reductions in muscle mass and contractile force, we leveraged matched transcriptomic, mass, and force measurements from individual muscles for mediation analysis. Given the high dimensionality of transcriptomic data, mediation analysis at the individual gene level was not feasible. To address this, we first applied gene set variation analysis (GSVA) (Hänzelmann et al. 2013), a single-sample gene set enrichment method, to derive enrichment scores for the 50 Hallmark gene sets and reduce dimensionality (Figure 5a) (Liberzon et al. 2015). However, despite the low redundancy among these curated gene sets, enrichment scores were strongly correlated across many gene sets (r > 0.8; Figure 5a), indicating shared underlying transcriptional activity.

**FIGURE 5.**
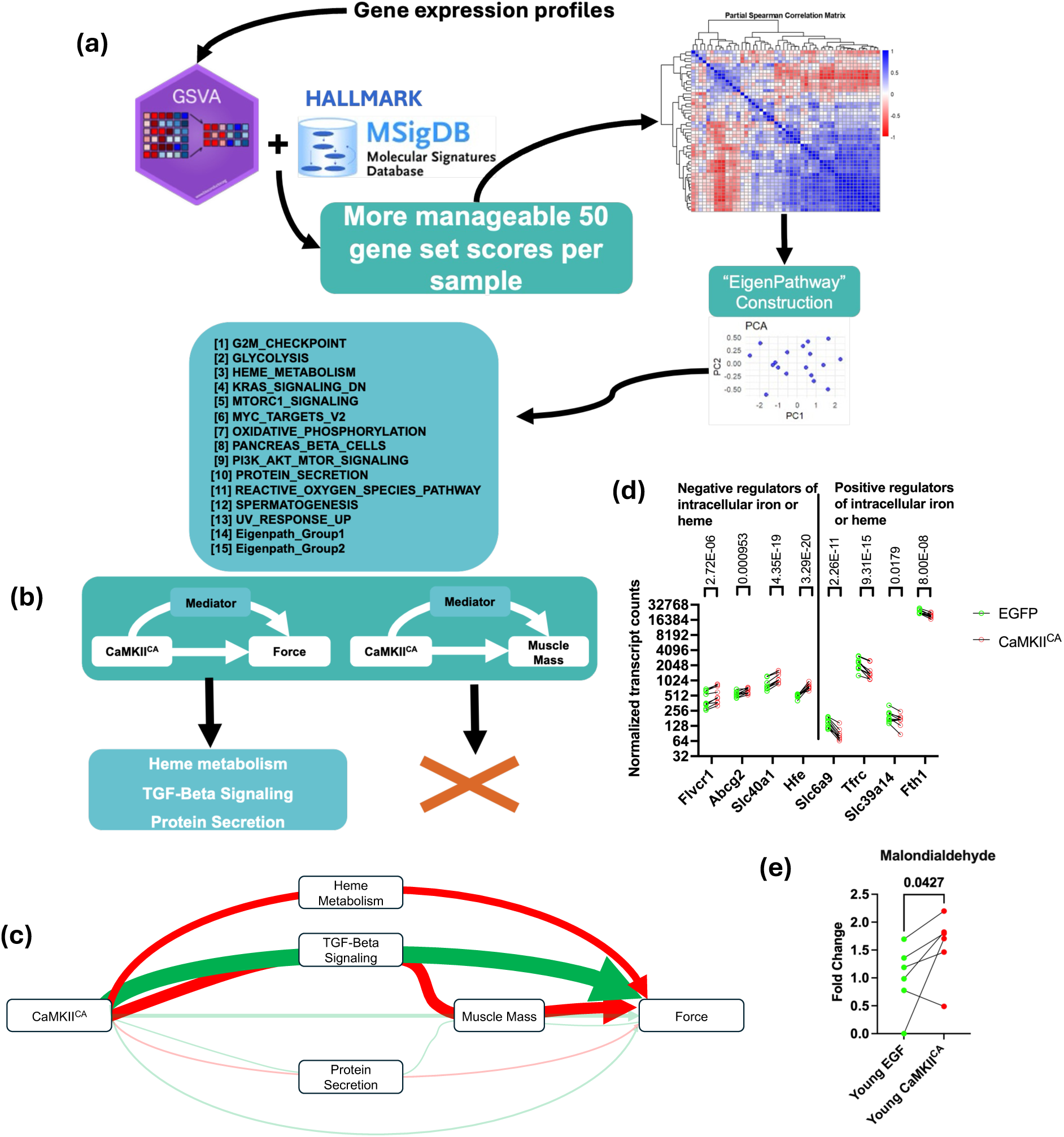
Gene set variation analysis (GSVA) and mediation analysis identify heme metabolism as a direct mediator of CaMKII^CA^-induced muscle weakness. (a) To identify potential mediating effects in the transcriptome, GSVA was used to derive enrichment scores for the 50 hallmark gene sets in each sample. Partial Spearman correlation analysis was performed on the gene set enrichment scores, adjusting for subject ID, to identify redundant gene sets, defined as those with | ρ| ≥ 0.8, with two clusters identified and reduced to “eigenpathways” based on principal component analysis with the first principal component representing the new “eigenpathway” score. (b) All potential mediators were evaluated through two mediation models, one mediating contractile force and another mediating muscle mass, using the “mediation” package in R. After correcting for multiple comparisons, only three significant mediators were identified. (c) The Diagram of the full structural equation model (SEM) model. Each arrow represents a path from CaMKII^CA^ treatment (independent variable) to contractile force (outcome), with boxes representing mediators of paths. The color and size of the arrows denote effect (red negative, green positive, and size magnitude), and the opaque arrows indicate significant paths. (d) Normalized transcript counts of heme and iron regulators. Whole transcriptome multiple-test adjusted p-values from DESeq2 analysis are included in the figure. (e) Malondialdehyde measured by mass spectrometry in CaMKII^CA^- and EGFP-expressing TA muscles. n = 6 per condition, one-tailed t-test, p-value in graph.

To minimize redundancy, we aggregated highly correlated gene sets (r > 0.8) into composite variables termed “eigenpathways.” This yielded 15 candidate gene sets, 13 from the original Hallmark collection and two composite eigenpathways, for mediation analysis. Each was evaluated as a potential mediator of CaMKII^CA^’s effects on muscle mass and contractile force. No gene set significantly mediated the effect on muscle mass following correction for multiple comparisons. In contrast, three gene sets, heme metabolism, TGF-β signaling, and protein secretion, significantly mediated the reduction in contractile force (Figure 5b).

To integrate these findings, we constructed a structural equation model (SEM) in which heme metabolism, TGF-β signaling, and protein secretion were entered as parallel mediators of CaMKII^CA^ effects on muscle mass and contractile force. The initial model allowed each gene set to have only a direct path to force, independent of muscle mass. Indirect paths through muscle mass were subsequently included when the modification index exceeded 3.84, indicating a statistically significant improvement in model fit. In the final SEM, heme metabolism retained a single direct path to force, whereas TGF-β signaling and protein secretion showed both direct and muscle mass-mediated paths to force (Figure 5c). TGF-β signaling exhibited an inconsistent mediation pattern, where a significant negative indirect effect on force via reduced muscle mass was partially offset by a smaller positive direct effect. Protein secretion did not retain significant mediation effects in the full model.

Collectively, these results identify heme metabolism as the principal transcriptomic mediator of CaMKII^CA^-induced contractile impairment, independent of muscle mass loss. Given the central role of the heme metabolism pathway in mediating CaMKII^CA^-induced muscle weakness and its close relationship with iron homeostasis (Dutt et al. 2022), we examined the expression of key genes related to heme and iron regulation in the RNA-seq data. We found that CaMKII^CA^ caused significant changes in transcript abundance in both pathways. Specifically, negative regulators of intracellular heme and iron were significantly upregulated, including heme exporters Flvcr1 and Abcg2, the iron exporter Slc40a1 (Ferroportin), and Hfe (Homeostatic Iron Regulator). Conversely, positive regulators of intracellular heme and iron were significantly downregulated, including Slc6a9 (GlyT1 glycine transporter) that imports glycine for heme synthesis, iron importers Tfrc (transferrin receptor 1) and Slc39a14 (Zip14), as well as the iron storage protein Fth1 (ferritin heavy chain) (Figure 5d). This expression pattern is consistent with a negative feedback response to intracellular iron overload (Galy et al. 2024).

Iron excess is known to promote oxidative stress by catalyzing the production of hydroxyl radicals, which damage DNA, proteins, and lipids (Galy et al. 2024). The activation of the long-patch base-excision DNA repair pathway (Figure 4d) and a significant elevation of lipid peroxidation product malondialdehyde in CaMKII^CA^-expressing muscle as measured by mass spectrometry (Figure 5e) support the concept that chronic CaMKII activation promotes oxidative stress, potentially by inducing iron overload.

### Inhibiting CaMKII in aged muscles improves contractility without inducing hypertrophy

To test whether the transcriptomic rejuvenation induced by CN19o translates into functional benefit, we injected AAV9-tMCK-CN19o in one TA of 21-month-old mice and AAV9-tMCK-EGFP in the contralateral TA. Five weeks later, CN19o-expressing muscles generated higher absolute force across the 1–150 Hz stimulation range (linear mixed-effect model, estimate = 0.45 [0.2, 0.7], p = 5.04 × 10^-4^, Figure 6a). This functional enhancement occurred despite a trend towards reduced muscle mass in the CN19o-treated TAs (∼5% reduction vs. EGFP, p = 0.0725; Figure 6b). Consequently, when force was normalized to muscle mass, CN19o-expressing muscles demonstrated a pronounced improvement (linear mixed-effect model, estimate = 0.79 [0.42, 1.17], p =5.09 × 10^-5^, Figure 6c). Mediation analysis confirmed a significant direct positive effect of CN19o on force (Average Direct Effect [ADE]: 0.64, p < 2 × 10^-16^, Figure 6d). While the slight reduction in muscle mass exerted a minor counteracting indirect effect (Average Causal Mediation Effect [ACME]: -0.194, p = 0.004), the combined outcome was a significant net increase in contractile force driven by CN19o (Total Effect: 0.45, p < 2 × 10^-16^). H&E staining showed a similar gross appearance of CN19o-expressing muscle fibers to EGFP controls, with no signs of endomysial inflammation or necrotic fibers in either condition (Figure 6e). The median minimum Feret diameters of individual CN19o-expressing muscles were not significantly different from contralateral EGFP-expressing controls (Figure 6f, p = 0.2437), but when the measurements of fibers were pooled by treatments, CN19o-expressing group showed significantly smaller minimum Feret diameters than the EGFP group (Figure 6g, p < 0.0001), and a leftward shift in the distribution of minimum Feret diameters (Figure 6h). These observations were consistent with mild atrophy induced by CN19o.

**FIGURE 6.**
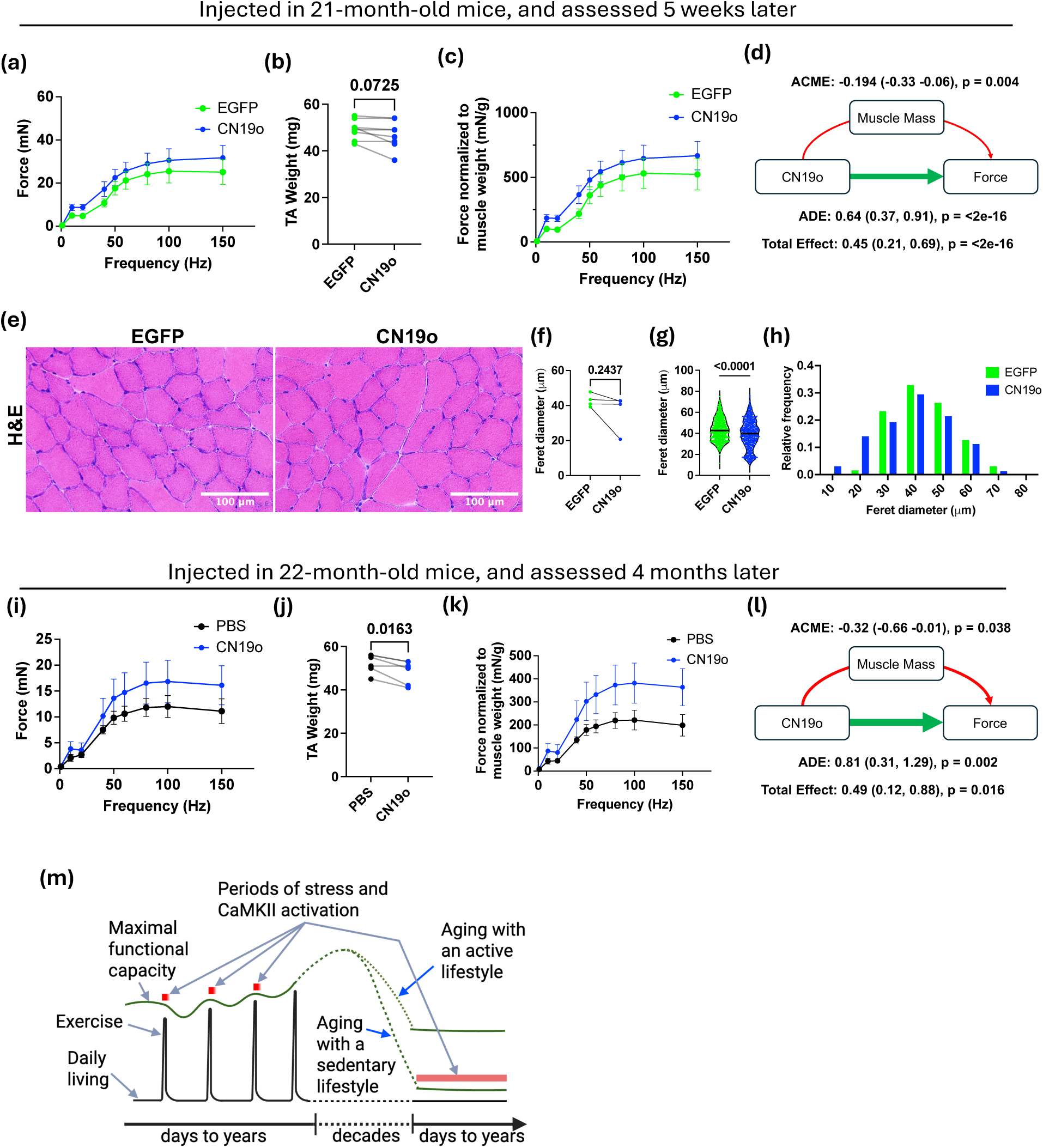
Inhibiting CaMKII in aged muscles improves contractility without inducing hypertrophy. (a) In vivo isometric force of TA muscles in response to peroneal nerve stimulation, 5 weeks post-injection of AAV-tMCK-CN19o (one TA) or AAV9-tMCK-EGFP (contralateral TA); n = 8 per condition; linear mixed-effects model, estimate = 0.45 (0.2, 0.7), p = 5.04 × 10^-4^. (b) TA muscle mass 5 weeks post-injection. n = 8 pairs; paired Student’s t-test, p-value in graph. (c) Isometric force normalized to TA mass. n = 8 per condition; linear mixed-effects model, estimate = 0.79 (0.42, 1.17), p = 5.09 × 10^-5^. (d) Mediation analysis (mediation” R package). (e) Representative H&E staining of TA cross-sections 6 weeks post-injection. (f) Muscle fiber minimum Feret diameters. Each point represents the median of ∼200 fibers per muscle. n = 4 muscles per treatment; paired Student’s t-test, p-value in graph. (g) Muscle fiber minimum Feret diameters, pooled by treatments. n = 839 (EGFP), n = 882 (CN19o), Mann-Whitney test, p-value in graph. (h) Histogram of relative frequency distributions of muscle fiber minimum Feret diameters. n = 839 (EGFP), n = 882 (CN19o). (i) In vivo isometric force of TA muscles in response to peroneal nerve stimulation, 4 months post-injection of AAV-tMCK-CN19o (one TA) or PBS (contralateral TA); n = 6 per condition; linear mixed-effects model, estimate = 0.5 (0.11, 0.89), p = 0.0131. (j) TA muscle mass 4 months post-injection. n = 6 pairs; paired Student’s t-test, p-value in graph. (k) Isometric force normalized to TA mass. n = 6 per condition; linear mixed-effects model, estimate = 0.96 (0.33, 1.59), p = 0.0033. (l) Mediation analysis. (m) Conceptual model. Green lines and green dotted lines indicate maximal functional capacity; the black curve indicates functional demand from daily living and exercise; red blocks indicate stress-induced CaMKII activation. See Discussion for details.

To exclude the possibility that the apparent benefit of CN19o was merely an artifact of EGFP-induced force suppression in the contralateral control muscles (Agbulut et al. 2006), a separate cohort of 22-month-old mice received AAV9-tMCK-CN19o in one TA and PBS in the contralateral TA. Four months later, CN19o-expressing muscles again generated significantly greater absolute force (linear mixed-effect model, estimate = 0.5 [0.11, 0.89], p = 0.0131, Figure 6i) and normalized force (linear mixed-effect model, estimate = 0.96 [0.33, 1.59], p =0.0033, Figure 6k) despite a modest decrease in muscle mass (∼7%, p = 0.0163; Figure 6j). Mediation analysis confirmed a robust direct effect of CN19o on force (ADE: 0.81, p = 0.002) that was only partially offset by reduced mass (ACME: -0.32, p = 0.038), yielding a significant net force increase (Total Effect: 0.49, p = 0.016; Figure 6l).

## 3 DISCUSSION

Excessive ROS and dysregulated intracellular Ca^2+^ signaling are well-established drivers of muscle dysfunction (Meng & Yu 2010; Michelucci et al. 2021). Paradoxically, both are also essential mediators of exercise-induced muscle adaptation, the most effective intervention against sarcopenia (Egan & Sharples 2023). We show that CaMKII, a key transducer of ROS and Ca^2+^ signals during exercise, is upregulated and persistently activated in aged skeletal muscle. Inhibition of CaMKII in aged muscle partially restores muscle strength and youthful transcriptional profiles, whereas constitutive activation of CaMKII in young muscle induces features of muscle aging at the functional, morphological, and molecular scales. Our results identify chronic CaMKII activation as a causal contributor to muscle dysfunction in aging, and support that CaMKII exemplifies the antagonistic pleiotropy theory of aging (Wang et al. 2021), which posits that traits harmful in older organisms are favored by natural selection if they confer benefits in at a younger age (Rose 1994, pp.62–78).

To model sustained CaMKII activation of aging muscle, we used intramuscular AAV9 delivery to express a constitutively active CaMKII mutant (CaMKII^CA^) under the control of the muscle-specific tMCK promoter in young muscle. Although CaMKII^CA^ expression reduced endogenous CaMKII levels and autophosphorylation at T287, overall CaMKII activity was elevated, as confirmed by the CaMKII activity reporter CaMKAR. This treatment led to a reduction in muscle force that initially occurred independently of the reduction in muscle mass. CaMKII^CA^ also induced the formation of tubular aggregates, a feature frequently observed in older animals (Boncompagni et al. 2012; Boncompagni et al. 2021), and progressive mitochondrial disorganization. By 9 months post-transduction, muscle atrophy contributed significantly to strength deficits, as supported by mediation analysis. This temporal pattern of functional decline preceding loss of muscle mass recapitulates a hallmark feature of human muscle aging and satisfies the definition of sarcopenia (Sayer & Cruz-Jentoft 2022). A recent study using intramuscular AAV1 delivery of constitutively active CaMKIIβ under the broadly active CMV promoter similarly reported reduced muscle mass and force in young mice over a 3-month time frame (Eguchi & Yamanashi 2022). However, that study did not examine aging muscle or assess histological or transcriptional outcomes. In contrast, our muscle-specific models incorporate these analyses to elucidate how sustained CaMKII activity contributes to muscle aging and provide evidence for the benefits of CaMKII inhibition in aged muscles.

RNA-seq analysis revealed that young CaMKII^CA^-expressing muscles exhibit a transcriptomic profile closely resembling that of aged muscle. These findings support the pleiotropic nature of CaMKII signaling. To elucidate the mechanisms by which chronic CaMKII activation drives muscle aging and functional decline, we first performed GSEA and identified several pathways similarly activated by CaMKII^CA^ and aging, including stress response, proteasome, and inflammatory pathways. Notably, partial inhibition of CaMKII by CN19o in aged muscle subtly but globally reversed age-associated transcriptional changes, supporting CaMKII activation as a driver of the aging-associated gene expression program. However, while chronic inflammation is a well-established contributor to sarcopenia (Granic et al. 2023), and CaMKII is known to activate NF-κB (Singh et al. 2009; Martin et al. 2018; Yao et al. 2022), NF-κB blockade did not ameliorate CaMKII^CA^-induced functional decline, indicating that additional CaMKII-dependent pathways are involved.

Mediation analysis identified the enrichment score of the heme metabolism gene set in GSVA as a mediator of CaMKII^CA^-induced force decline. Consistent with the close relationship between heme and iron homeostasis, our RNA-seq data show coordinated changes in heme- and iron-related transcripts: genes promoting heme and iron accumulation were downregulated, while those restricting their intracellular levels were upregulated. Several of these genes are regulated by negative feedback mechanisms (Furuyama et al. 2007; Dutt et al. 2022), and therefore, their expression changes suggest a compensatory response to intracellular heme and iron excess in CaMKII^CA^-expressing muscles. Iron overload is cytotoxic and has been implicated in age-related skeletal muscle dysfunction by promoting oxidative stress, cellular senescence, and ferroptosis (Terrell et al. 2023; Alves et al. 2023; Galy et al. 2024). CaMKII^CA^-expressing muscle showed increased activation of stress response and long-patch base excision DNA repair, and elevated levels of the lipid peroxidation product malondialdehyde, consistent with excessive iron-mediated oxidative stress. Because CaMKII is activated by oxidation, sustained CaMKII activity and heme/iron dysregulation may form a self-amplifying vicious cycle in aging muscle. Mechanistically, because alteration of heme and iron-related genes was observed in CaMKII^CA^-expressing muscles relative to the contralateral control muscles of the same mice, we speculate that CaMKII^CA^ influences heme and iron homeostasis in a tissue-restricted, rather than systemic fashion. Ongoing studies will investigate how excessive CaMKII activation leads to heme/iron overload and assess its relationship to the aging-related decline of muscle function, iron overload, mitochondrial disorganization, and tubular aggregate formation.

Our data reveal a deleterious role for CaMKII in aging skeletal muscle, which contrasts with its established beneficial role in exercise-induced adaptation. This contrast highlights the importance of the duration, intensity, and context of CaMKII activation. We propose that both exercise-induced and aging-associated CaMKII activation are triggered by disruption of cellular homeostasis (Figure 6m). In young muscle, such disruption typically occurs only during deliberate exercise, where CaMKII activation is transient, proportional to physiological demand, and resolved with recovery (Rose & Hargreaves 2003; Rose et al. 2006; Egan et al. 2010). Aging, by reducing maximal functional capacity, lowers the threshold for homeostatic disruption, such that activities of daily living may chronically activate CaMKII and render it maladaptive. Our model is consistent with the recommendation to use exercise as an intervention for sarcopenia, as it preserves functional capacity and may reduce aberrant CaMKII activation.

For individuals unable to engage in effective training, pharmacological CaMKII inhibition may provide a complementary strategy. We propose that temporally targeted CaMKII inhibition during resting periods without interfering with its adaptive function in exercise could optimize functional outcomes. The FDA-approved JAK inhibitor ruxolitinib was recently identified as a potent CaMKII inhibitor at therapeutic concentrations (Reyes Gaido et al. 2023). Notably, ruxolitinib has been shown to enhance physical performance in naturally aged and progeria mouse models, as well as to increase muscle mass in patients with myelofibrosis (Xu et al. 2015; Griveau et al. 2020; Lucijanic et al. 2021). Although these effects were attributed to JAK inhibition, further studies are warranted to evaluate the contribution of CaMKII inhibition to these benefits.

### Limitations

We used the AAV9-tMCK vector to specifically deliver CaMKII^CA^ and CN19o to muscle fibers in young and old mice, enabling paired comparisons and mediation analyses. While AAV9 is minimally immunogenic, and AAV9-tMCK-EGFP was used as a control, vector-related effects cannot be completely ruled out. As CaMKII^CA^ is based on the γ isoform, and skeletal muscle expresses additional isoforms with different localization and functions, it may not fully recapitulate aging-associated dysregulation of endogenous CaMKII. Additionally, the low expression of CN19o, as indicated by RNA-seq, may have limited the extent of CaMKII inhibition. Finally, to minimize animal usage, control experiments were conducted on the contralateral limb; however, this approach does not rule out potential cross-communication between limbs injected with different AAV9 vectors via myokines or other signaling molecules. Future studies will address these limitations and utilize pharmacological agents and genetic models to more precisely manipulate CaMKII and assess its translational relevance in human muscle.

## 4 EXPERIMENTAL PROCEDURES

### 4.1 Animals

An equal number of male and female C57BL/6J mice were obtained from the National Institute on Aging (NIA) Aged Rodent Colony. All animal procedures adhered to National Institutes of Health (NIH) guidelines and were approved by the Johns Hopkins University School of Medicine Institutional Animal Care and Use Committee. Mice were housed in the Johns Hopkins animal facility under a 12-hour light/dark cycle and provided 2018 Teklad Global 18% Protein Rodent Diet ad libitum. The housing environment was maintained at 22 ± 1°C with a humidity level of 40 ± 10%.

### 4.2 Western blotting

Muscle samples were homogenized in T-PER™ reagent (Thermo Fisher Scientific, #78510) containing cOmplete™ Protease Inhibitor Cocktail (Roche, #11697498001) and PhosSTOP™ Phosphatase Inhibitor Cocktail (Roche, #4906845001) using a Precellys homogenizer. After centrifugation at 10,000 × g for 10 minutes, 5 μg (for mitochondrial proteins) or 20 μg (for all other proteins) of protein from supernatants was resolved on NuPAGE Tris-Bis gels and transferred to PVDF membranes. Total protein was visualized using Pierce Reversible Protein Stain (Thermo Fisher Scientific, 24585) before membranes were blocked with EveryBlot buffer (Bio-Rad, 12010020) for 5 minutes at room temperature. Membranes were incubated with primary antibodies at optimized dilutions, followed by HRP-conjugated secondary antibodies (20 ng/mL). Immunoreactive bands were detected using SuperSignal™ West Dura Substrate (Thermo Fisher Scientific) on a Bio-Rad Gel Doc XRS+ system and analyzed with Image Lab software (v6.1), with normalization to total protein content as determined by reversible staining.

Primary antibodies: total-CaMKII (BD biosciences, 611292, 1:3000), pT287-CaMKII (Cell Signaling Technology, D21E4, 1:2000), FLAG (Thermo Fisher Scientific, MA1-91878, 1:2000), EGFP (Abcam, ab290, 1:2000), OxPhos Rodent WB Antibody Cocktail (Thermo Fisher Scientific, 458099, 1:1000).

### 4.3 AAV9 viral vector construction and injection

The AAV9-tMCK-CaMKII^CA^ construct was generated by appending a triple-FLAG tag (MDYKDHDGDYKDHDIDYKDDDDK) and a flexible linker (GGGGS) to the N-terminus of mouse CaMK2g (NM_001039138) coding sequence. To create the constitutively active CaMKII^CA^, the threonine residue at position 287 was mutated to aspartic acid (T287D) through codon substitution to GAC. This sequence was subcloned into a pAAV-tMCK-WPRE vector backbone, and the recombinant AAV9-tMCK-CaMKII^CA^ viral particles were produced by Vector BioLabs (Great Valley Parkway, Malvern PA).

AAV9-tMCK-CN19o was developed using a synthetic DNA sequence (ATGTACCCTTACGACGTACCGGACTACGCTGGCGGGGGAGGGAGTAAGCGGGCTCCAAAACTCGGCCAGATCGGAAGACAGAAAGCCGTGGATATTGAAGATTGA) encoding an HA tag followed by a flexible linker (GGGGS) and the CN19o peptide (KRAPKLGQIGRQKAVDIED). This construct was similarly cloned into the pAAV-tMCK-WPRE vector backbone, with viral packaging performed by Vector BioLabs.

The IκB super repressor (ISR) virus, AAV9-tMCK-ISR, was generated by introducing S32A and S36A mutations into the mouse Nfkbia coding sequence (NM_010907.2). The modified cDNA was then cloned into the pAAV[Exp]-tMCK-WPRE vector and packaged into AAV9 by VectorBuilder (Chicago, IL).

The control virus AAV9-tMCK-eGFP (SKU VB5037) was obtained from Vector BioLabs.

The viral constructs were diluted in sterile phosphate-buffered saline (PBS) to a final concentration of 2.0 × 10^12^ viral genomes/mL and administered using 31-gauge needles. Each tibialis anterior (TA) muscle of anesthetized mice (2% isoflurane) received two 25 μL injections at separate sites, delivering a total dose of 1.0 × 10^11^ viral genomes per muscle (Cho et al. 2019). To minimize experimental bias, control and treatment viruses were alternated between left and right legs across different animals, and the experimenter conducting the muscle contractility assay was blinded to the identity of the injected viruses.

Isolated muscle fiber CaMKAR imaging from skeletal muscle injected with AAV9-tMCK-CaMKII^CA^. All the animal procedures and protocols were reviewed and approved by the Institutional Animal Care and Use Committees of the University of Maryland. Male C57BL/6J mice (Charles River, Wilmington, MA) were used. All mice used were 12 weeks of age. Environmental conditions were maintained with a 12-h light/dark cycle and constant temperature (21–23°C) and humidity (55 ± 10%). The cages contained corncob bedding (Harlan Teklad 7902) and environmental enrichment (cotton nestlet). Mice were supplied with dry chow (irradiated rodent diet; Harlan Teklad 2981) and allowed water ad libitum. C57BL/6 mice were infected at 8 weeks of age via intramuscular injection into the flexor digitorum brevis (FDB) muscle. A total dose of 1.0 × 10¹¹ viral genomes per muscle was administered. Subsequently, to facilitate the expression of the CaMKAR reporter system in vivo, electroporation was performed on the FDB muscles of 11-week-old mice (Banks et al. 2021). Subcutaneous injection of 20-30 μl of hyaluronidase (2.5 mg/ml) was administered into the plantar pad of a mouse anesthetized with 3% - 4.5% isoflurane in O₂ (1 L/min), using a 33-gauge needle. One hour later, the mouse was again anesthetized, and ∼60-70 μg of CaMKAR plasmid DNA was injected into the footpad. The animal was kept on an isothermal pad for 5 minutes; then, two surgical stainless-steel electrodes were placed subcutaneously close to the proximal and distal tendons of the FDB muscle and 20 pulses of 100 V/cm, 20 ms in duration, were applied at 1 Hz via a commercial high current capacity output stage (ECM 830, BTX, Harvard Apparatus, Holliston, MA). One week later, single muscle fibers were enzymatically dissociated from the injected FDB muscles and cultured as described below (Bibollet et al. 2023). FDB muscle was isolated from male adult mice, enzymatically dissociated with collagenase type I (Sigma-Aldrich, St. Louis, MO) in spinner minimum essential medium (S-MEM; Cat. No. 11380-037; Gibco, Carlsbad, CA) with 10% FBS (Cat. #100–106; Gemini Bio-Products, West Sacramento, CA) for 3-4 hours at 37°C. Muscle was then gently triturated to separate fibers in S-MEM with FBS 10%. Fibers were plated in MEM culture media with 2% FBS on glass-bottomed dishes (MatTek Cor, Ashland, MA, Cat. No. P35G-1.0-14-C) coated with laminin (Thermo Fisher, Rockford, IL, Cat. No. 23017-015). Fibers were maintained in culture for 2 to 20 hours at 37 °C, 5% CO2 before the experiments. Positively transfected fibers were identified using an intensiometric approach by assessing the fluorescence expression profile excited with 488 nm and emitted light collected with a long-pass filter >510 nm. Cultured skeletal muscle FDB fibers expressing CaMKAR constructs were imaged in 2 mL of L-15 media (ionic composition in mM: 137 NaCl, 5.7 KCl, 1.26 CaCl2, 1.8 MgCl2, pH 7.4; Life Technologies, Carlsbad, CA). All single fiber recordings were performed at room temperature, 22°C. Individual muscle fibers expressing CaMKAR were excited with a 488 nm laser, and the fluorescence emitted >505 nm was acquired with an Olympus IX70 inverted microscope equipped with an Olympus FLUOVIEW 500 laser scanning confocal microscope imaging system using an Olympus 40X UPlanXApo objective. Fluorescence was calculated by the average of the intracellular fluorescent signal, considering three different regions of the isolated fibers (after subtracting the background extracellular signal). Five to seven single fibers per culture plate were analyzed independently for each animal. Regions of interest (ROIs) in the images were quantified using ImageJ software (National Institutes of Health, Bethesda, MD).

### 4.3 In vivo muscle contractility measurements

Muscle contractility was assessed using the Aurora 1300A system, as previously described (Westbrook et al. 2020). Briefly, mice were anesthetized with 4% isoflurane for induction and maintained at 1.5% during the procedure. They were positioned supine on a warming pad to sustain body temperature, with the knee stabilized and the foot secured to the footplate. Percutaneous stimulation of the peroneal nerve was applied using 100 μs pulses, with current optimized to elicit maximal isometric force. The force-frequency relationship was assessed using 250 ms pulse trains ranging from 1 to 150 Hz. Data were exported from DMA-HT analysis software (Aurora Scientific), analyzed in R, and visualized using GraphPad Prism 10.

### 4.4 Histology

Whole TA muscles were frozen in liquid nitrogen-cooled isopentane and sectioned into 8 μm sections. The samples were stained with the Hematoxylin and Eosin staining kit (Vector Laboratories, H-3502). Protocols for modified Gomori’s trichrome staining, SDH, and COX staining were obtained from the Washington University Neuromuscular Disease Center.

For TEM, samples were fixed in 2.5% glutaraldehyde, 3 mM MgCl_2_ in 0.1 M sodium cacodylate buffer, pH 7.2, overnight at 4 °C. After buffer rinse, samples were postfixed in 2% osmium tetroxide in 0.1 M sodium cacodylate for at least one hour (no more than two) on ice in the dark and samples rinsed in distilled water, dehydrated in a graded series of ethanol and embedded in Epon (PolySci) resin. Samples were polymerized at 60 °C overnight.

Thin sections, 60 to 90 nm, were cut with a diamond knife on a Leica UCT ultramicrotome and picked up with 2x1 mm Formvar copper slot grids. Grids were stained with 3% uranyl acetate (aq.) followed by lead citrate and observed with a Hitachi 7600 TEM at 80 kV. Images were captured with an AMT CCD XR80 (8-megapixel camera - side mount AMT XR80 – high-resolution high-speed camera).

Light and TEM images were analyzed in Fiji ImageJ with treatments blinded to the analyzer (Schindelin et al. 2012). For light microscopy, for each muscle sample, two locations within a single mid-belly cross-section were imaged, with one image taken from the superficial region and the other from the deep region. All muscle fibers in both images were measured for their minimum Feret diameter, also termed the lesser diameter, a robust index of muscle fiber size minimally affected by the plane of section (Dubowitz et al. 2020). For each muscle, 200–300 fibers were measured, which is sufficient to reliably quantify muscle fiber size (Dubowitz et al. 2020).

Mitochondrial area was quantified by manually outlining all mitochondria in randomly selected, non-overlapping longitudinal muscle section images using Fiji, followed by area measurement. Four muscle samples per treatment group were analyzed.

### 4.5 RNA sequencing and analyses

Total RNA was isolated from TA muscle samples five weeks after viral injection using the Direct-zol RNA MiniPrep Kit (Zymo Research, R2050). RNA quality was assessed with a Fragment Analyzer (Advanced Analytical Technologies, Inc.), and all samples had an RNA Quality Number (RQN) >7.0. mRNA enrichment was performed using 1 μg of total RNA per sample with the NEBNext® Poly(A) mRNA Magnetic Isolation Module (New England Biolabs, E7490L), followed by RNA library preparation with the NEBNext® Ultra^TM^ II Directional RNA Library Prep Kit (New England Biolabs, E7765L). Each library was uniquely barcoded using the NEBNext Multiplex Oligos for Illumina (96 Unique Dual Index Primer Pairs, New England Biolabs, E6440S). The RNA libraries were then submitted to the Johns Hopkins Genetic Resources Core Facility and sequenced on a NovaSeq X Plus (Illumina, Inc.) to generate 2 × 150 bp paired-end reads.

Transcriptomic data collected by RNA sequencing (RNA-seq) was analyzed to determine the genes that are present in each sample and condition, their expression levels, and the differences between expression levels among experiment conditions, as follows. Sequencing reads were mapped to the mouse genome version GRCm39 with the spliced alignment tool STAR 2.7.10a (Dobin et al. 2013). The aligned reads were assembled with PsiCLASS v.1.0.3 (Song et al. 2019) to create gene and transcript models. Transcripts were then assigned to known genes from the GENCODE v.M28 reference gene set augmented with the three transgene sequences (EGFP, CN19o, CaMKIICA) as additional “chromosomes”. Lastly, DESeq2 (Anders & Huber 2010) was used to quantify the expression levels and determine differentially expressed genes. Additional visualizations, including volcano plots and plots of principal component analysis (PCA) components, were visualized with custom R scripts (R v.4.1.3).

To account for similarities as the sequence level between the CaMKIICA transgene and the endogenous Camk2g mouse gene, which contribute to multi-mapped reads that can alter the expression calculations for the two genes, we performed two analyses. The first analysis used the PsiCLASS gene annotations as created above. The second analysis identified and distinguished ‘shared’ versus ‘unique’ areas of the two genes, with the rationale that unique areas will provide a more accurate assessment of the genes’ expression levels. For the Camk2g gene, we separated the ‘unique’ 3’ most exon (chr14:20784942-20786920; internal gene ID ‘chr14.111872.e1’) from the rest of the gene, which shows sequence similarity to the transgene. For the CamKIICA transgene, we separated the first 200 bp (internal gene ID: ‘chr_CaMKIICA.402115’), which includes the 84 bp linear sequence, as ‘unique’, from the rest of the gene. These unique gene portions can be used to more accurately assess and compare the expression levels of these two genes. All the other genes’ expression levels are unaffected by the change in annotation.

Gene Set Enrichment Analysis was performed using GSEA 4.4.0 with pre-ranked gene lists according to Log2(Fold Change) of Old EGFP vs. Young EGFP, CaMKII^CA^ vs. Young EGFP, and CN19o vs. Old EGFP (Subramanian et al. 2005). The hallmark gene sets were first used to understand the broad biological states of the transcriptomes (Liberzon et al. 2015), followed by more detailed analyses using the curated canonic pathways (M2.CP.Reactome). The gene sets that had FDR q-value <0.25 were imported into Cytoscape (v3.10.3) to generate Enrichment maps to reduce the redundancy in gene sets, using the EnrichmentMap App (v3.5.0) (Merico et al. 2010).

Correlation analysis was performed in R using the “cor.test” function. For each comparison at the transcript level, transcripts that were significant by p < 0.05 in either condition were included. For visualization purposes, BH-corrected Fisher’s combined p-values were used for bubble size.

Log₂-transformed expression matrices were subjected to gene-set variation analysis (GSVA) in R (“GSVA” version 1.48.0, “GSEABase” version 1.60.0) using the “hallmarks” gene sets. Partial Spearman correlations controlling for subject ID were computed with pcor.test (“ppcor” version 1.1), and an igraph network was constructed from edges with |ρ| ≥ 0.8; each connected component containing > 1 variable was collapsed to its first principal component (“stats” version 4.4.1), termed an “eigenpathway.”

### 4.6 Contractile Force Analysis

To assess changes in contractile force between conditions, linear mixed-effects models were generated in R using the “lmer” function of the “lme4” package (version 1.1.35.5). The “boxcox” function of the “MASS” package (version 7.3.61) was used to transform the force readings to ensure that the assumption of linearity and normality was met. Additionally, frequency values were log-transformed to ensure linearity, resulting in a final model of: BoxCoxForce ∼ Condition+log(Frequency)+(1|ID), with ID representing random effects of subject to account for within-subject and repeated measures design.

### 4.7 Mediation Analysis

To determine the role of muscle mass in CaMKII^CA^- and CN19o-induced changes to contractile force, we employed mediation analysis using the “mediation” package (version 4.5.0). Mediator models were constructed as: muscle ∼ Condition + (1|ID), and outcome models as: BoxCoxForce ∼ Condition + muscle + log(Frequency) + (1|ID). For the construction of the combined SEM model (“lavaan” version 0.6.19), four mediators were included: heme metabolism, TGF-β signaling, protein secretion, and muscle mass, with sequential mediation paths from TGF-β signaling and protein secretion to muscle mass based on modification indices and biological plausibility.

### 4.8 Metabolomics analysis

The malondialdehyde measures were derived from an ongoing metabolomic study.

#### Sample Preparation

Muscle tissues from all experimental groups were processed in parallel using a standardized extraction workflow. The precisely weighed samples were subjected to homogenization in a solvent mixture composed of 99% LC-MS-grade acetonitrile and 1% formic acid, using a multi-probe sonicator to ensure efficient homogenization. The homogenates were then transferred to extraction plates equipped with sorbent filters specifically designed to precipitate proteins and remove phospholipids. The metabolite solution was passed through the sorbent filter using a Positive Pressure-96 processor, and the purified metabolite solution was collected in an MS plate, while simultaneously retaining proteins and phospholipids in the sorbent filter. The extracted solution were then concentrated to a final volume of 150 µL in preparation for mass spectrometry.

#### Data Acquisition

Mass spectrometry was performed on a Thermo Scientific IQ-X platform interfaced with a Vanquish UHPLC system. Reversed-phase separation was achieved using a Discovery HSF5 column (Sigma-Aldrich), operated under a 9-minute gradient—7 minutes for metabolite separation and 2 minutes for re-equilibration. The autosampler maintained samples at 4°C throughout, with 2 µL injected per run, and the column maintained at 35°C. The mobile phase consisted of MS-grade water and acetonitrile, both containing 0.1% formic acid.

In addition, Hydrophilic Interaction Liquid Chromatography (HILIC) was performed using an ACQUITY Premier BEH Amide column paired with a VanGuard FIT Cartridge (Waters). This method employed a 15-minute runtime (13.5 minutes for data capture and 1.5 minutes for re-equilibration). The mobile phase system consisted of 95:5 20 mM ammonium acetate with 20 mM ammonium hydroxide (pH 9) in water and 100% acetonitrile. Column temperature and sample conditions mirrored those used for reversed-phase analysis.

Prior to sample acquisition, system calibration was conducted to ensure optimal sensitivity and mass accuracy. Metabolite identities were assigned using accurate mass. Peak areas were integrated from the chromatograms and normalized to tissue mass to obtain final relative metabolite concentrations.

#### Data Analysis

Downstream analysis was carried out using Gigantest’s Laboratory Information Management System (LIMS) with additional custom pipelines written in Python and R. Chromatographic peaks were initially processed using machine learning-based algorithms, followed by manual inspection to confirm correct peak boundaries and integrations.

## Author Contributions

Q.W. and M.B. wrote the manuscript. Q.W. and P.A. carried out project administration and supervision. Q.W., M.B., T.H.C., E.H.O., and P.A. designed experiments. M.B., Q.W., T.C., G. R.-S., E.H.-O., C.A., L.F., A.L, Q.-L. X, and A.H. analyzed data. M.B., Q.W., A.L., W.A.F., G.R.-S. S.J.J. performed experiments. All authors discussed and edited the manuscript.

## Acknowledgement

1. M. B. is supported by the National Institute on Aging (2T32AG058527-06A1) and the National Institute of Arthritis and Musculoskeletal and Skin Diseases (5T32AR048522-20). Q.W. is supported by the Glenn Foundation for Medical Research and AFAR Grants for Junior Faculty, Karen and Ethan Leder CIM Human Aging Project Scholarship, Nathan Shock scholarship, the Johns Hopkins Older Americans Independence Center Research Education Core grant (NIA P30AG021334), and the Johns Hopkins Center for AIDS Research Faculty Development Grant P30AI094189. E.H.-O. was supported by the National Institutes of Health grant R01-AR075726. L.F. is supported by NIH award R35GM156374.

We thank Dr. Jeremy Walston for facilitating this project, Dr. Rafael deCabo for providing the 33-month-old mice, Landon Bechdel for assisting with experiments, Barbara Smith of the Johns Hopkins Microscope Facility for TEM, Emily Elassal for assisting with TEM data quantification, and Dr. Mark E. Anderson for critical reading and suggestions.

## Data Availability Statement

Source data are available upon reasonable request. The bulk RNA-sequencing data have been deposited as GSE****** upon the publication of the manuscript.

---

**SUPPLEMENTARY FIGURE 1.**
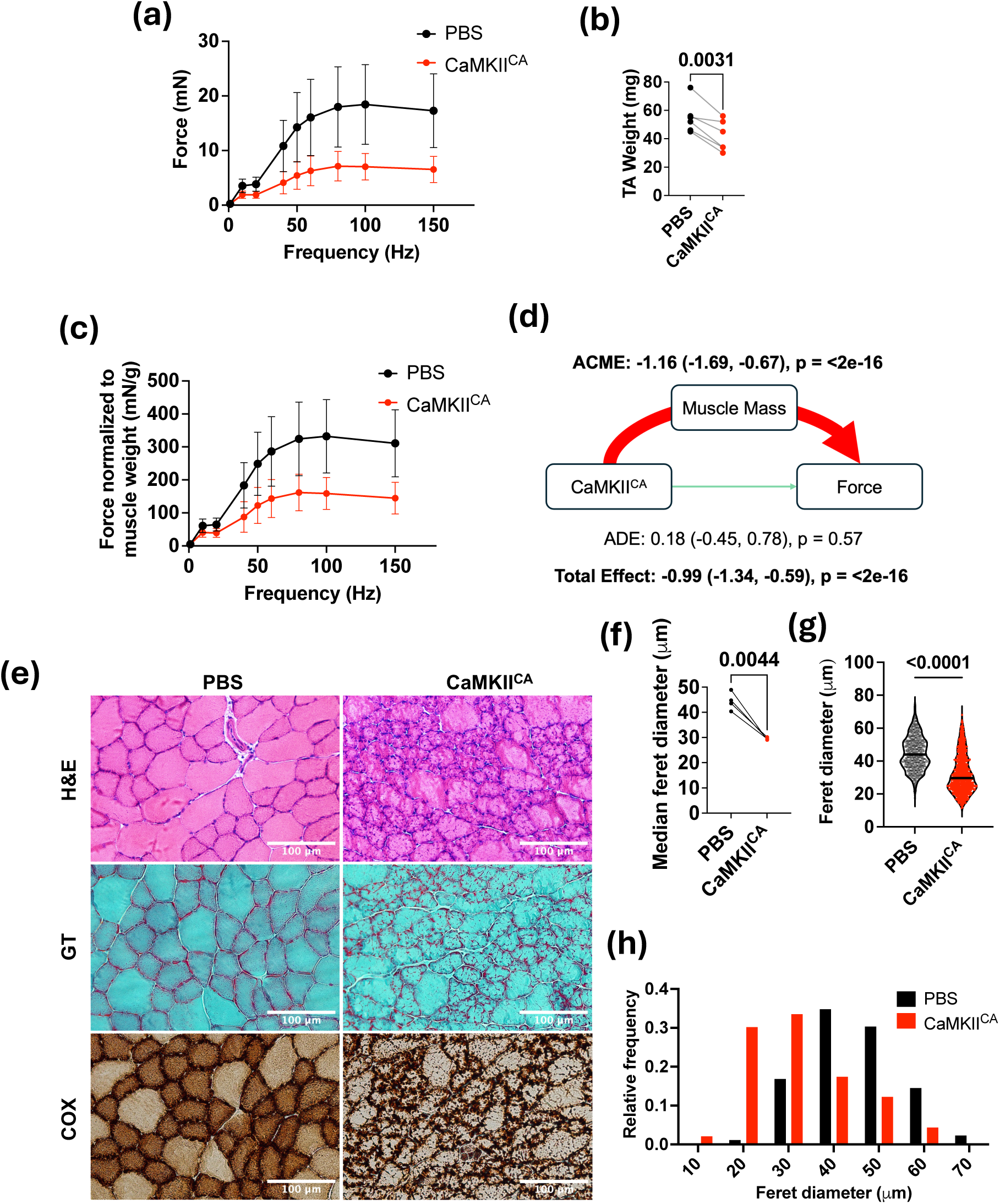
Chronic CaMKII activation further impairs muscle contractility, mass, and histological structures. (a) In vivo isometric force of TA muscles in response to peroneal nerve stimulation at different frequencies, 8.5 months post-injection of PBS (one TA) or AAV9-tMCK-CaMKII^CA^ (contralateral TA). n = 6 per condition; linear mixed-effects model, estimate = - 0.99 (-1.39,-0.59), p = 4.21 × 10^-6^. (b) TA muscle mass 8.8 months post-injection. n = 6 pairs; paired Student’s t-test, p-value in graph. (c) Isometric force normalized to TA mass. n = 5; linear mixed-effects model, estimate = -1.25 (-1.87, -0.62), p = 1.48 × 10^-4^. (d) Mediation analysis (“mediation” R package). (e) Representative H&E, modified Gomori’s Trichrome (GT), and COX staining of TA cross-sections 8.8 months post-injection. (f) Quantification of muscle fiber minimum Feret diameters. Each point represents the median of ∼200–300 fibers per muscle. n = 4 muscles per treatment; paired Student’s t-test, p-value in graph. (g) Muscle fiber minimum Feret diameters, pooled by treatment. n = 778 (PBS), n = 1,337 (CaMKII^CA^). (h) Histogram of relative frequency distributions of muscle fiber minimum Feret diameters. n = 778 (PBS), n = 1,337 (CaMKII^CA^).

**SUPPLEMENTARY FIGURE 2.**
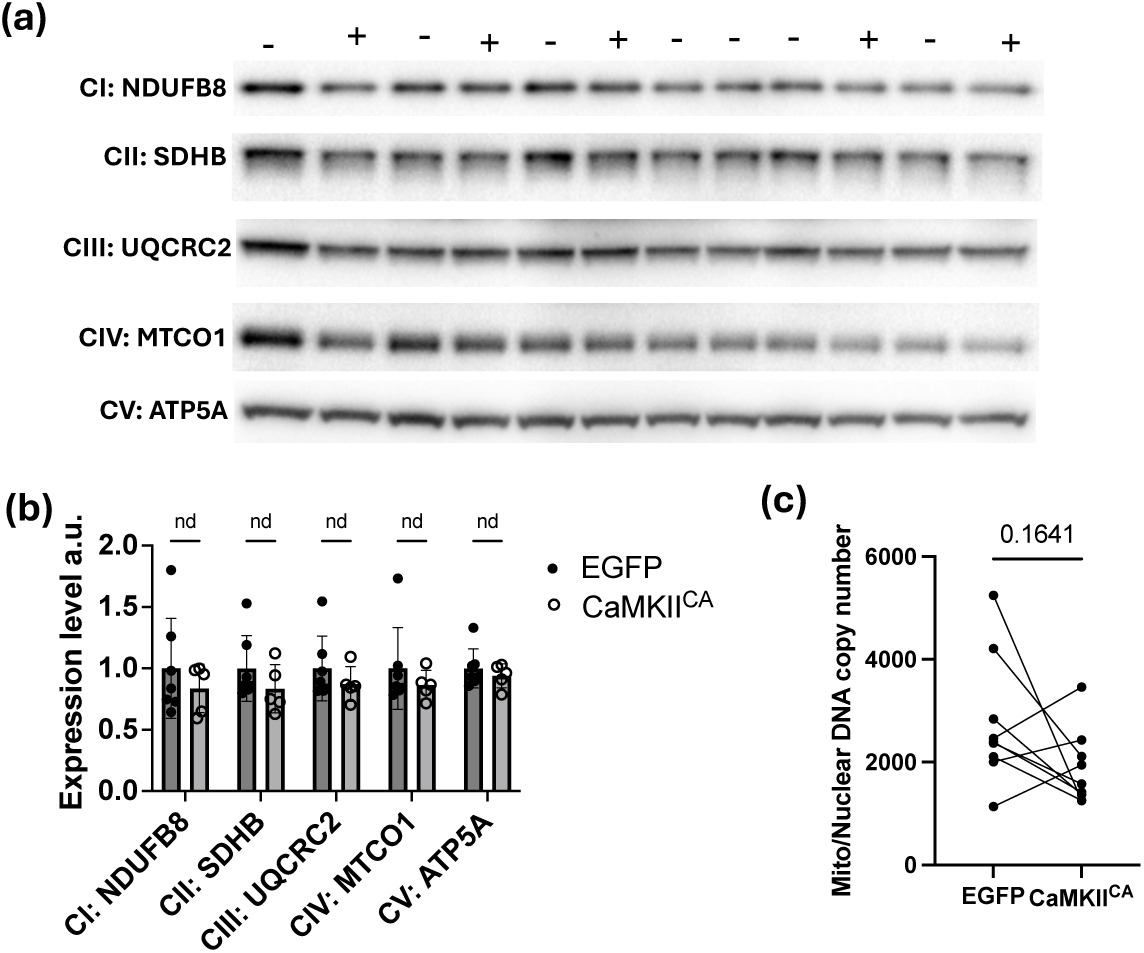
Short-term CaMKII^CA^ expression does not alter mitochondrial complex abundance or mtDNA copy number in TA muscles 7 weeks post-injection. (a) Western blot detection of representative subunits of mitochondrial complexes I–V. (+) denotes AAV9-CaMKII^CA^-injected TA, and (-) denotes AAV9-EGFP-injected TA. (b) Quantification of individual subunit expression levels relative to total protein staining. n = 7 (EGFP), n = 5 (CaMKII^CA^); multiple t-tests, nd = not a discovery (not significant). (c) Quantification of mitochondrial DNA to nuclear DNA copy number ratio by qPCR. n = 8 pairs, paired Student’s t-test, p-value in graph.

**SUPPLEMENTARY FIGURE 3.**
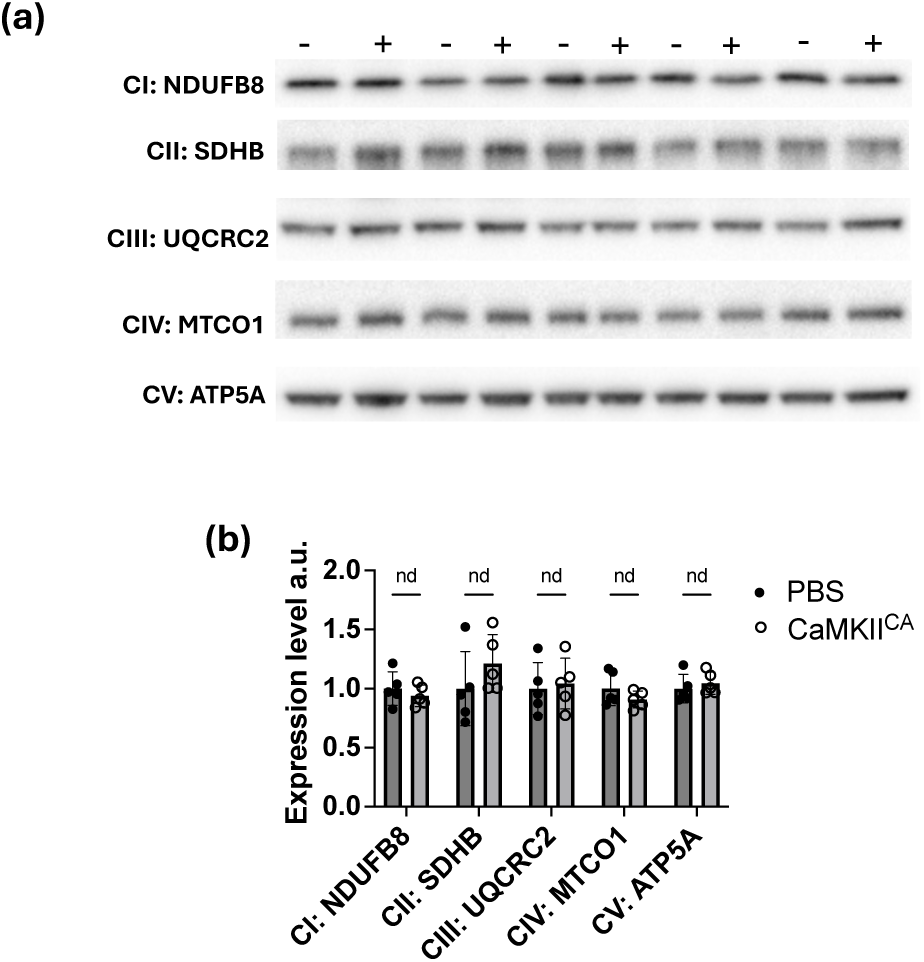
Long-term CaMKII^CA^ expression does not change the abundance of mitochondrial complexes I–V (C–CV) in TA muscles 9 months post-injection. (a). Western blot detecting the representative subunits of mitochondrial complexes I–V. (+) denotes AAV9-CaMKII^CA^-injected TA, and (-) denotes PBS-injected TA. (b) Quantification of individual subunit expression levels. n = 5 pairs; paired multiple t-tests, nd = not a discovery (not significant).

